# Stress resilience is an active and multifactorial process manifested by structural, functional, and molecular changes in synapses

**DOI:** 10.1101/2022.05.19.492644

**Authors:** E. Bączyńska, M. Zaręba-Kozioł, B. Ruszczycki, A. Krzystyniak, T. Wójtowicz, K. Bijata, B. Pochwat, M. Magnowska, M. Roszkowska, I. Figiel, J. Masternak, A. Pytyś, J. Dzwonek, R. Worch, K.H. Olszyński, A.D. Wardak, P. Szymczak, J. Labus, K. Radwańska, P. Jahołkowski, A. Hogendorf, E. Ponimaskin, R.K. Filipkowski, B. Szewczyk, M. Bijata, J Włodarczyk

## Abstract

Stress resilience is the ability of neuronal networks to maintain their function despite the stress exposure. Using a mouse model we here investigate stress resilience phenomenon. To assess the resilient and anhedonic behavioral phenotypes developed after the induction of chronic unpredictable stress, we quantitatively characterized the structural and functional plasticity of excitatory synapses in the hippocampus using a combination of proteomic, electrophysiological, and imaging methods. Our results indicate that stress resilience is an active and multifactorial process manifested by structural, functional, and molecular changes in synapses. We reveal that chronic stress influences palmitoylation of synaptic proteins, whose profiles differ between resilient and anhedonic animals. The changes in palmitoylation are predominantly related with the glutamate receptor signaling thus affects synaptic transmission and associated structures of dendritic spines. We show that stress resilience is associated with structural compensatory plasticity of the postsynaptic parts of synapses in CA1 subfield of the hippocampus.

**One Sentence Summary:** Compensatory remodeling of dendritic spines at the structural and molecular levels underlies stress resilience.

## INTRODUCTION

The resilience phenomenon has been broadly investigated in physics^1–3^, geoscience^4^, botany^5^, ecology^6^, sociology^7,8^, economics^9^, and neuroscience^10,11^. The definition of resilience is not universal and is often determined by researchers based on their own scientific interests^10,12–15^. The common feature of resilience manifests in its dynamic nature, i.e., the ability of a system (e.g., a material, a plant, or a brain) exposed to a harmful stimulus to absorb, accommodate or adapt to the effects of the stress in an efficient manner by adjusting its structure, organization, physiological mechanisms, metabolic regulation, mode of action, etc. The resilience phenomenon is also commonly observed in human neurological diseases, particularly in neurodegenerative and neuropsychiatric diseases^11,16–20^. Despite exposure to traumatic events or even to the presence of advanced pathophysiological changes, some people do not exhibit behavioral and psychological symptoms. The origin of such a diverse response to stressful conditions at the level of brain plasticity is an unexplored field of modern neuropsychiatry and neuroscience. Many researchers point to the principal role of genetic predispositions in stress resilience, while others see its origin in protective factors, reward system, brain reserve or aberrant functions of key molecules^21–27^. Despite the broad research in this field, the impact of the environmental factors contributing to stress resilience is still largely unexplored. One of the fundamental questions that still lacks a clear answer is whether the stress resilience observed in adulthood might be an actively regulated process influenced by environmental factors^28^. Several findings indicate that stress resilience can be pharmacologically enhanced upon treatment with glutamate receptor antagonists, e.g., ketamine^26,29–32^. However, the changes in the molecular landscape underlying drug-induced stress resilience have not yet been identified in the treatment of stress-related disorders.

Major depressive disorder (MDD) is one of the commonly occurring forms of stress-related disorders and has been extensively studied both in humans and in animal models^33,34^. The etiology of MDD is complex and idiopathic. Many factors influence the development of MDD, including genetic predispositions and the environment^35,36^. Nevertheless, chronic stress is a key contributor to MDD development due to its impact on the dysregulation of the hypothalamic–pituitary–adrenal axis and adrenal steroids^37,38^. Observations of the diverse distribution of glucocorticoid receptors within brain regions reveals the medial prefrontal cortex, hippocampus, and amygdala as the structures most affected by chronic stress in MDD patients and animals displaying depressive-like behavior^38–40^. Of importance, the broad, interdisciplinary characterization of the hippocampus enables the integration of molecular, functional, and imaging data in the context of animal behavior. Moreover, *postmortem* studies of MDD subjects revealed that antidepressive action requires synaptic plasticity in the hippocampus^41,42^. Multiple studies postulate that the mechanisms of MDD are described by the hypotheses of the monoamine-, inflammation-, neurotrophy-, circadian- or GABA/glutamate mediated hypotheses^43,44^. However, none of them is convincing enough to explain the molecular mechanism or the combination of all of the symptoms and subtypes of MDD^43,44^. Nevertheless, all these hypotheses share a common concept that aberrant synaptic plasticity underlies depressive symptoms and that pathology-related structural changes are caused by the increased excitatory transmission induced by chronic stress^38,45–48^. Furthermore, the restoration of excitatory structural connectivity in the rodent hippocampus and prefrontal cortex with ketamine, a fast-acting NMDA receptor antagonist, is sufficient to return animals to a healthy behavioral state^49–52^.

The changes in structural connectivity are related to the processes occurring in the excitatory synapses^53^. Most of the excitatory synapses are located on dendritic spines, which are small, motile membrane protrusions of neurons^54^. Dendritic spine remodeling involves alterations in spine morphology and/or density^53^. The structural plasticity of dendritic spines is a hallmark of physiological (learning and memory) and pathological (neurological and neuropsychiatric disorders) conditions^55–57^. Studies of depressive-like behavior in animal models revealed abnormalities in the maintenance of the strength of synaptic connections that were manifested by a loss of spines and/or an increase in the proportion of immature, thin forms of spines. This finding suggests that the mechanism responsible for the transformation of dendritic spines might be a key factor underlying depressive-like behaviors^57,58^. However, this phenomenon appears to be more complex due to the observations that dendritic spine remodeling is not necessary for the short-term antidepressant effect of ketamine, but it may be essential to sustain the remission of depressive-like behavior^51,52,59^. These studies point out the downstream signaling pathways underlying the regulation of dendritic spine shape and function as a possible decisive molecular target for promoting stress resilience and preventing pathogenesis.

There are no reports describing how the structure of dendritic spines and the underlying molecular landscape of the tetrapartite synapse are affected in resilient animals following chronic stress and how components of the tetrapartite synapse regulate this process. Recently, S-palmitoylation (S-PALM) has gained attention due to its role in stress-related neuropsychiatric diseases^60^ and the remodeling of dendritic spines^61,62^. S-PALM is a fast-acting posttranslational lipid modification in which palmitoyl group is attached to cysteine residues in peptides and proteins. S-PALM is one of the most unique posttranslational modifications because, unlike other lipid modifications, S-PALM is reversible^63,64^ and may be regulated by the neuronal activity triggered by environmental factors^65^ or pharmacological treatment^62^. S-PALM controls protein stability, receptor trafficking, and protein–protein interactions and thus contributes to synaptic plasticity, e.g., modulation of long-term potentiation (LTP)^61,65–67^. However, the role of S-PALM in stress resilience and the contribution of S-PALM to the dendritic spine remodeling is still an open subject and requires further study.

In the present study, we address the following questions: *i)* Is stress resilience a process that is actively developed in the adult brain?; *ii)* To what extent does chronic stress affect the molecular architecture of the synapse in resilient and anhedonic animals?; *iii)* Are the differences between resilient and anhedonic animals related to the structural and/or functional plasticity of excitatory synapses?; *iv)* Is it possible to explain the potential differences in underlying synaptic plasticity between resilient and anhedonic animals in terms of the palmitoylation of synaptic proteins?

To answer the aforementioned questions, we applied chronic unpredictable stress (CUS) leading to the development of resilient and anhedonic behavior. The evaluation of the functional, structural, and molecular readouts of excitatory synaptic plasticity in the hippocampus, in relation to animal behavior, was performed by a multidisciplinary and quantitative methodological approach that include mass spectrometry, electrophysiology, and fluorescent confocal imaging. The presented results indicate that stress resilience is a multifactorial phenomenon that actively develops during adulthood. Moreover, we demonstrate that the developed stress-resilient state is associated with aberrant synaptic plasticity at the structural, functional and molecular levels.

## RESULTS

### Chronic stress leads to the development of stress resilience and anhedonia in adult mice

To investigate whether stress resilience might develop during adulthood, we employed CUS to induce a behavioral stress response in adult animals. Ten-week-old male C57BL6J mice were subjected to 2-weeks of chronic and unpredictable stress in which different types of stressors (restraint stress, social defeat stress, tail suspension stress, predator-induced stress) applied in a pseudorandom manner on consecutive days within the dark and light phases of the 12/12 cycle (see *Materials and Methods* for details). We demonstrated that the implementation of such an intense CUS procedure leads to the development of anhedonic and resilient behaviors, as determined by the sucrose preference test (SPT, Figure 1A-C). Anhedonic mice were considered by sucrose preference following chronic stress (SPT1) < 70.7% (defined by the difference between the control and stressed groups being higher than the two standard deviations i.e., >2xSD, established in our recent studies^68^). In turn, mice were considered stress resilient after CUS when they did not exhibit anhedonia in SPT1 (sucrose preference > 70.7%). After CUS, approximately 50% of mice developed anhedonic behavior, and the rest were classified as stress-resilient (Figure 1C)^68,69^. Moreover, mice subjected to CUS (both anhedonic and resilient) exhibited less robust body weight gain than that of the non-stressed control animals. Thus, body weight is a physiological indicator of exposure to chronic stress (Figure 1D)^70,71^. Taken together, our data indicate that implementation of the designed CUS procedure leads to the synchronic development of anhedonic and resilient behavior in genetically homogenous adult mice, constituting a promising animal model of stress resilience. Figure 1E-G shows the behavioral parameters (SPT, FST) and body weight gain of the mice that were used in experiments described in the next parts of manuscript. We observed a significant negative correlation between the SPT and FST results (Figure S1, p=0.02).

**Figure 1.**
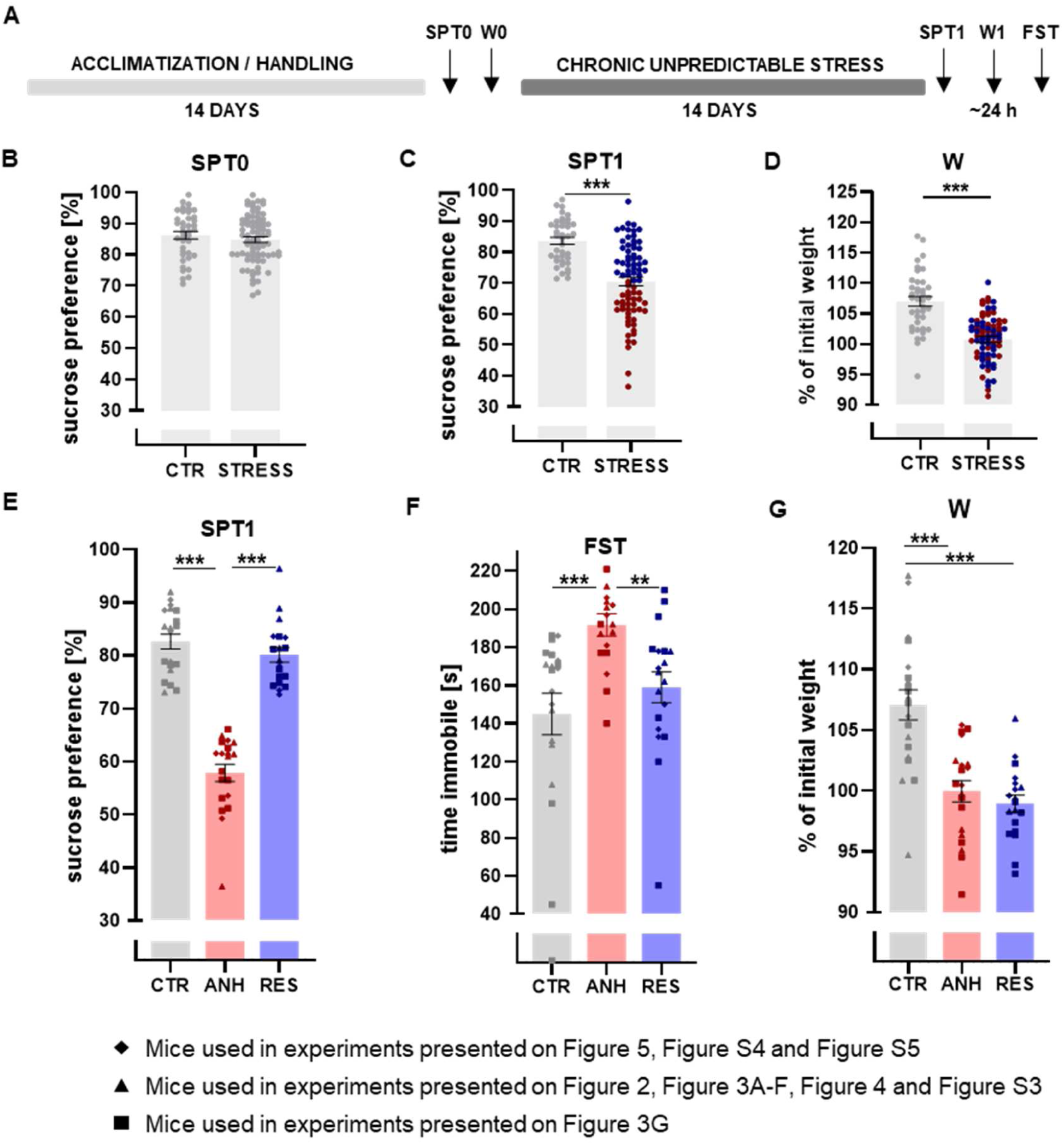
Behavioral evaluation of depressive-like behavior of anhedonic (ANH) and resilient (RES) animals following chronic unpredictable stress (CUS) as well as control animals (CTR). **(A)** Schematic view of the experimental design of the CUS model. **(B)** Baseline sucrose preference test (SPT0). **(C)** Sucrose preference test after CUS (SPT1), anhedonic animals are marked in red and resilient animals in blue **(D)** Body weight gain after CUS (W) n_CTR_=39, n_STRESS_=78 **(E-G)** Behavioral parameters of animals used in experiments described in the next figures n=18-20. **(E)** Sucrose Preference Test after CUS (SPT1). **(F)** Forced swim test after CUS (FST). **(G)** Body weight gain after CUS (W). The data are presented as the mean ± SEM. *p < 0.05; ***p < 0.001. (B-C – two tailed t-test, D - Mann Whitney test E-F-Dunn’s multiple comparisons test, G – Brown-Forsythe ANOVA test followed by Tamhane’s T2 multiple comparisons test.

### Chronic stress affects the expression of synaptic proteins in the hippocampus

As a first step to determine the molecular fingerprint of stress resilience, we employed a high-throughput proteomic approach using mass spectrometry to generate a comprehensive view of the *in vivo* proteins level in the hippocampal synaptoneurosomes of control, resilient and anhedonic animals. To isolate synaptoneurosomes we used a method based on ultracentrifugation and a density gradient^72,73^. We evaluated the quality of the synaptoneurosome preparation using electron microscopy (Figure S2). To identify differentially expressed proteins across behavioral groups, their protein levels were determined and comparatively analyzed in five biological replicates per group. Additionally, the clustering heatmaps of Pearson correlation coefficients (PCCs) of peptide signal intensity were determined using logarithm 2 transformed protein abundance data (Figure 2A) to evaluate the variability in the proteomic analysis within biological replicates and behavioral groups. The correlation matrix shows the PCC values for all experimental groups. The matrix values for each stressed group differed from the matrix values for the control group. Moreover, the variation in the matrix values for the control group was small, in contrast to the variations for the other two groups (Figure 2A). Volcano plots depict the changes in protein expression among behavioral groups (Figure 2B). The fold change logarithm (base 2) is on the x-axis, and the negative logarithm of the false discovery rate (p value) (base 10) is on the y-axis. Protein levels with p value < 0.05 and fold changes < -0.5 and > 0.5 were considered significantly different. We identified 6224 proteins with less than 1% FDR (false discovery rate). Our results revealed 44 differential synaptic proteins in resilient animals (24 upregulated and 20 downregulated) and 39 in the anhedonic group (20 upregulated and 19 downregulated) in comparison to non-stressed animals. The list of the proteins within the aforementioned groups is presented in Supplemental Tables 1A-B. Surprisingly, the levels of only 6 synaptic proteins significantly differed between the resilient and anhedonic groups: Ppp3ca, Recql, Tenm1, Fastkd1, Gcc2 and Zdhhc13 (one downregulated and five upregulated; for more details, see Supplemental Table 1C, Figure 2B). Altogether, chronic stress affected less than 1% of the identified synaptic proteins in the hippocampus. In KEGG pathway bioinformatics analysis we found that the most significantly enriched pathways in anhedonic mice were related to maintenance of postsynaptic specialization, postsynaptic density organization, structural constituent of postsynapse, receptor clustering, neurotransmitter receptor localization to postsynaptic specialization membrane and protein localization to postsynapse (Figure 2C). In resilient mice the most affected pathways were related to regulation of dendritic spines morphogenesis (Figure 2D).

**Figure 2.**
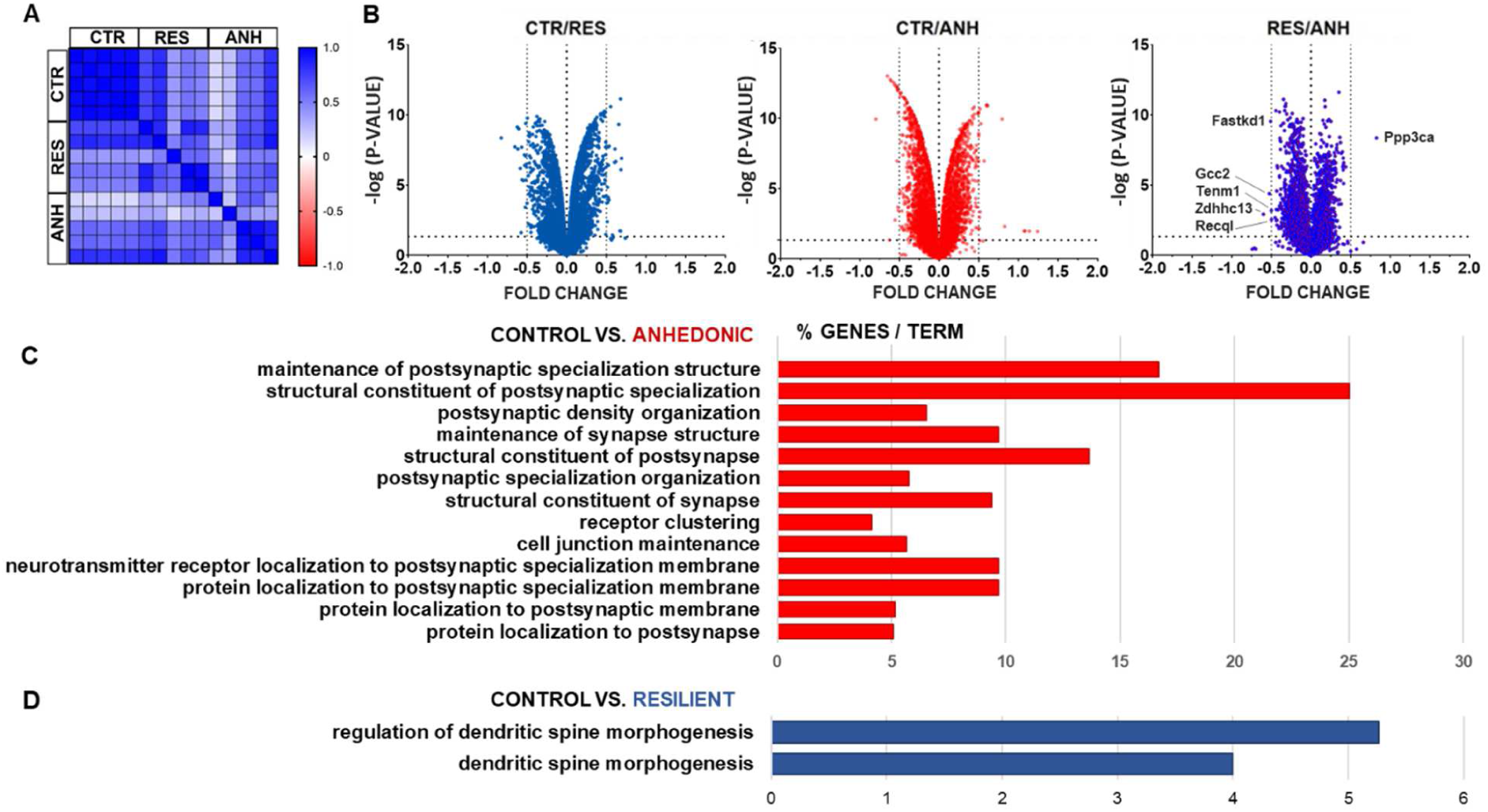
Analysis of differentially expressed synaptic proteins from proteomic profiling. **(A)** Matrix representation of Pearson correlation coefficients of protein abundances in 5 biological replicates. **(B)** Volcano plots representing changes in protein expression in CTR, ANH, and RES animals. The fold change log (base 2) is on the x-axis, and the negative false log discovery rate (p value, base 10) is on the y-axis. Negative fold change indicates increased expression while positive fold change decreased expression. (Two-stage linear step-up procedure of Benjamini, Krieger and Yekutieli). **(C-D)** Kyoto Encyclopedia of Genes and Genomes (KEGG) bioinformatics analysis of the significantly altered signaling pathways of the ANH and RES groups. The p value negative log (base 10) is on the x-axis.

### Distinctive palmitoylation of synaptic proteins in stress resilience

Due to the potential role of palmitoylation in chronic stress and neuropsychiatric diseases, our special attention was drawn to the palmitoyltransferase Zdhhc13 (DHHC22)^72,74^. We hypothesized that the altered level of palmitoyltransferases expression may alter the S-palmitoylation (S-PALM) profile of synaptic proteins differentially in stress resilient and anhedonic animals. We therefore took an unbiased proteomic approach based on the mass spectrometry PANIMoni method (biotin labeling of S-palmitoylated proteins)^72^ to identify proteins palmitoylated in response to the CUS procedure. We examined the level of similarity of protein content within the groups using principal component analysis (PCA) and clustering heatmaps of Pearson correlation coefficients (PCCs). We confirmed the reproducibility of the proteome preparation, enabling us to observe a global distinction between the control and stressed mice in the S-PALM profile of synaptic proteins. The results of the relative quantification experiment are summarized in Figure 3A-D. We identified S-PALM of 1199 synaptic proteins upon the CUS procedure in which 113 proteins were commonly changed in the same direction with a fold change < -1 and > 1 and p < 0.01 for all stressed mice, while 188 of synaptic protein differentiated the resilient phenotype from those of the anhedonic and control animals, constituting the S-PALM fingerprint of stress resilience (Supplemental Table 2A). To determine which cellular and synaptic processes were involved in stress resilience-induced turnover of palmitoylation, we analyzed the datasets of differentially palmitoylated proteins using the synaptic protein database ClueGO. In the ClueGO analysis, we identified functionally grouped networks linked to their GO biological processes and KEGG pathways, indicating palmitoylation-dependent biological processes that were associated mostly with the glutamate receptor signaling pathway (e.g., Daglb, Frrs1, Grin2b, Plcb1, Ptk2b, Shank3, Crtc1, Epha4, Rab3a,Tbc1d24), dendritic spine morphogenesis (e.g., Epha4, Kif1a, Shank3, Tanc2, Ptk2b, Tbc1d24, Erc1, Rab3a), receptor clustering (e.g., Grin2b, Lrp4, Shank3), synaptic vesicle turnover and localization (e.g., Ap3d1, Ap3m2, Htt, Kif1a, Rab3a, Shroom2, Tanc2, Ykt6, Tbc1d24, Ppp3cb), dopamine secretion (e.g., Abat, Kcna2, Prkcb, Rab3a, Syt3), and behavioral fear response (e.g., Grin2b, Shank3, Vdac1, Figure 3E-F). In addition, to address whether S-PALM is differentially manifested in different subregions of the hippocampus, the total S-PALM levels of all proteins were determined using acyl-biotin exchange method (Figure 3G). Obtained results indicate that chronic stress leads to decrease in S-PALM level mainly in the CA1 subregion of the hippocampus in comparison to the CA3 and dentate gyrus (DG) subregions.

**Figure 3.**
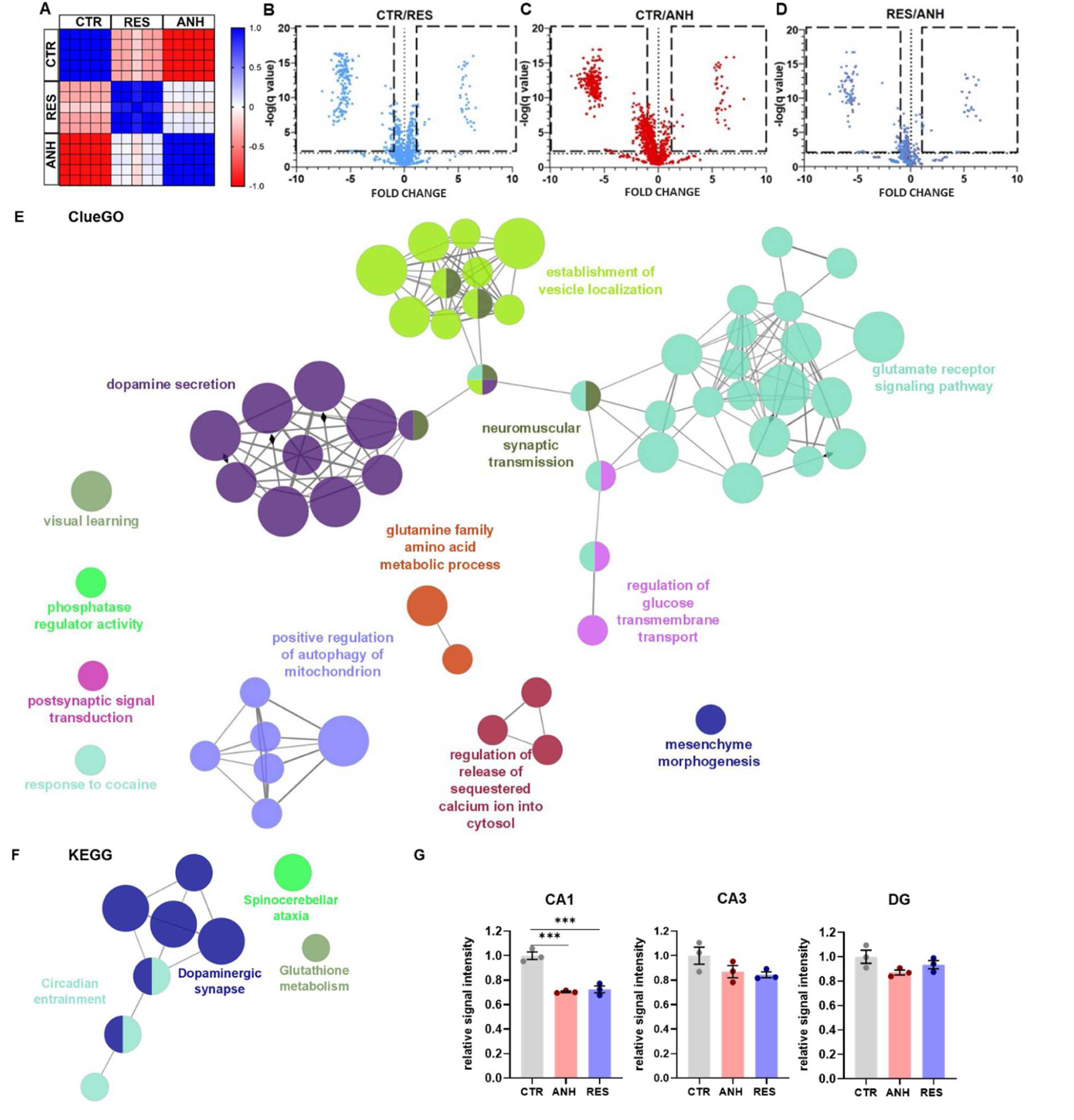
Analysis of differentially S-palmitoylated synaptic proteins from proteomic profiling. **(A)** Matrix representation of Pearson correlation coefficients of protein abundances in 5 biological replicates. **(B-D)** Volcano plots represent changes in protein S-palmitoylation in hippocampal synaptoneurosomes of CTR, ANH, and RES animals. The fold change log (base 2) is on the x-axis, and the negative false log discovery rate (p value) (base 10) is on the y-axis (Two-stage linear step-up procedure of Benjamini, Krieger and Yekutieli). **(E)** ClueGO, and **(F)** KEGG bioinformatics analysis of S-PALM proteins characteristic of stress resilience. Functionally grouped networks are linked to their GO biological processes and KEGG pathways. Each circle (node) represents a biological term consisting of various related proteins/genes. The node size represents the enrichment significance. Terms that belong to the same pathway are marked with the same color, and terms associated with two different pathways are marked with two colors. The connectivity (edges) between the terms in the network is derived from the kappa score (indicates the similarity of associated genes shared by different terms). Thicker edges indicate stronger similarity. Diamonds represent directed edges that link parent terms to child terms. Only the name of the most significant term in each group is shown to reduce the overlay. **(G)** Total palmitoylation of proteins in different subregions of the hippocampus (CA1, CA3, DG) of CTR, ANH, and RES animals. The experiment was performed in triplicate, using pooled samples from three independent animals per each condition (n = 9) The data are presented as the mean ± SEM; **p < 0.01; ***p < 0.001 (one-way ANOVA followed by Sidak’s multiple comparisons test).

### Chronic stress impairs postsynaptic glutamatergic neurotransmission in the hippocampus

Since we have observed changes in the palmitoylation profile of proteins related with the glutamate receptor signaling pathway in the hippocampus after chronic stress in resilient animals, we studied synaptic transmission by recording field excitatory postsynaptic potentials (fEPSPs) in acute hippocampal brain slices. To determine the α-amino-3-hydroxy-5-methyl-4-isoxazolepropionic acid receptors (AMPARs) and *N*-methyl-D-aspartate receptors (NMDARs) that contributed to synaptic transmission, we analyzed fEPSPs evoked in response to monotonically increased stimuli in the CA3-CA1 hippocampal projections in magnesium-free artificial cerebrospinal fluid (aCSF, Figure S3A-B)^75^. Experimental data were fitted with mathematical functions and compared for statistically significant differences by means of Monte Carlo randomization (see Materials and Methods, for details). Input–output curves in CA1 region showed that the amplitudes of the AMPAR-mediated fEPSPs as wells as the area of the NMDAR-mediated component of fEPSPs were not significantly different between groups (Figure S3C-D). We next studied the CA3 region and recorded responses of associational-commisural synapses in response to retrograde stimulation of the CA3 subregion^76^. We found that similar to CA1, CUS in the CA3 region resulted in no significant alteration of fEPSP amplitudes (Figure S4A). In contrast, the area of the NMDAR-mediated component of fEPSPs in CA3 region was significantly upregulated in ANH compared to RES group (p=0.031) while other comparison did not reveal significant differences (Monte-Carlo simulations, Figure S4). Thus, CUS resulted in differential modulation of glutamatergic AMPAR- and NMDAR-mediated synaptic responses in hippocampal subregions.

**Figure 4.**
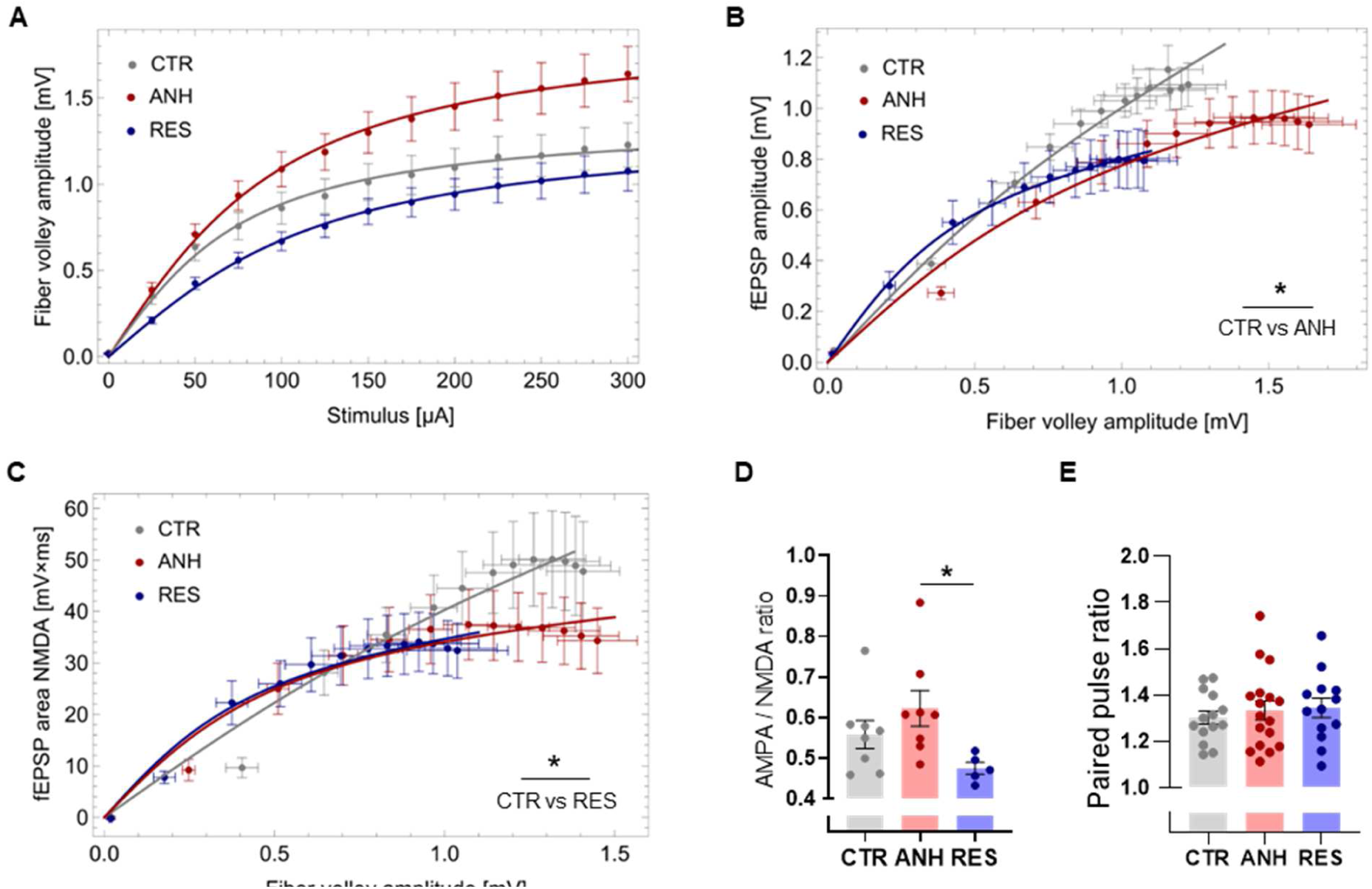
Chronic stress promotes differential changes in glutamatergic neurotransmission in ANH and RES hippocampi. **(A-C)** Average input–output relationships for fEPSPs recorded in the CA1 subregion of acute hippocampal slices and evoked in response to monotonically increased current stimuli applied to Shaffer collaterals. **(A)** fiber volley amplitudes were not significantly different following CUS but tended to vary between animal groups(p>0.05, n_slices_=15-20, N_animals_=5-6 per group, experimental data were fitted with a mathematical function and compared for statistically significant differences by means of three-dimensional Monte Carlo simulations (see Materials and Methods, for details). Experimental data were fitted with the function y(x) = a arctgx/b; see Materials and Methods for details). **(B-C)** fEPSP amplitudes and fEPSP area values normalized to fiber volley amplitudes in the same recordings (n_slices_=12-20, N_animals_=5-6 per group). **(B)** The magnitude of AMPAR-mediated fEPSPs recorded for the same fiber volley amplitude was significantly lower in the ANH but not the RES compared to the CTR group (p=0.03 and p=0.25 respectively). There was no difference between ANH and RES groups (p=0.54). **(C)** the NMDAR-mediated fEPSPs were significantly lower in RES group compared to CTR (p=0.01) while ANH vs RES and ANH vs CTR did no differ (p=0.055 and p=0.61, respectively). (D) The ANH group exhibited a higher AMPAR/NMDAR ratio estimated after DNQX application compared to that of the RES group (n_slices_=5-8, N_animals_=4-6 per group, one-way ANOVA, see Materials and Methods for details). (E) CUS did not affect the paired-pulse facilitation ratio (n_slices_=13-17, N_animals_=5-6 per group). Data are presented as the mean ± SEM. *p < 0.05.

To have further insight into this phenomenon, we have studied more thoroughly CA1 excitatory synapses. Surprisingly, we observed the largest apparent difference in the amplitudes of the fiber volley, although these differences were not significantly different (p>0.05, Monte Carlo method, Fig. 4A). The large variance in average fiber volley amplitudes could reflect changes in presynaptic axon function and/or the number of available afferent fibers following chronic stress^77^. Therefore, we visualized the fEPSP amplitude and the area vs. respective fiber volley amplitudes preceding these synaptic responses (Figure 4B-C). We found that the magnitude of AMPAR-mediated fEPSPs recorded for the same fiber volley amplitude was significantly lower in the ANH but not the RES compared to the CTR group (Fig. 4B). In addition, the NMDAR-mediated fEPSPs were significantly lower in RES group compared to CTR (Figure 4C). Therefore, stimulation of the same number of presynaptic afferents yielded less efficient excitatory AMPAR-mediated synaptic drive in the anhedonic group and NMDAR-mediated synaptic transmission in resilient group.

Consequently, we investigated whether chronic stress is associated with an altered synaptic AMPAR/NMDAR ratio. To this end, we compared the sensitivity of compound fEPSPs to an AMPAR-specific antagonist (DNQX), as described previously^78^. We found a larger sensitivity to DNQX and thus lower AMPAR/NMDAR ratio in the resilient group than in the anhedonic group (Figure 4D). The potentiation of synaptic release in response to paired-pulse stimulation was similar among the investigated groups (Figure 4E). Therefore, an altered presynaptic release was not attributed to any behavioral phenotype. Taken together, chronic stress affects glutamatergic neurotransmission in the hippocampus differently in the anhedonic group (less efficient excitatory drive and an increased AMPAR/NMDAR ratio) than in the resilient group. Since the observed functional alterations in the CA1 subregion of the hippocampus were limited to the postsynaptic part of the excitatory synapses, we subsequently analyzed the structural features of dendritic spines.

### Stress resilience is associated with structural compensation in the hippocampus

To determine how chronic stress affects dendritic spine structure, we performed DiI staining on hippocampal slices obtained from the second hemisphere and visualized them by fluorescence confocal microscopy. In the analysis of the hippocampus, we observed that the resilient animals differed from the control animals with respect to dendritic spine morphology. The spine density of the resilient animals was unchanged, while in the anhedonic group, decreased spine density was observed (p < 0.0001, Figure 5A, E, Figure S5). The quantitative analysis of spine morphometry in the entire hippocampus showed a decreased *length-to-head-width* ratio and spine *area* in the resilient group than in the controls (p < 0.05, Figure 5B, E, Figure S5). Upon analysis of the individual subregions (i.e., CA1, CA3 and DG), we identified morphological changes unique to a given subregion. In the CA3 subregion, the spines of RES animals showed a lower length-to-head-width ratio compared to that of CTR animals. Such changes usually indicate the maturation of the dendritic spines, however a more detailed morphometric analysis revealed that the changes in the length-to-head-width ratio were mainly due to a decrease in the length of the spine, while the surface area and head width did not increase (as is usually the case with spine maturation), but it was reduced which may indicate pathological changes resulting from chronic stress (Figure 5C, E, Figure S5). However, the CA1 subregion caught our special attention. In this subregion, the spines of ANH animals showed a higher length-to-head-width ratio compared to CTR animals and RES animals. In parallel, the spines of RES animals had a larger head width than that of anhedonic animals, and there was a clear upward trend compared to CTR animals (p = 0.10) (Figure 5C, E, Figure S5). In the CA1 subregion pyramidal neurons develop two dendritic trees covered with dendritic spines - basal dendritic tree located in *Stratum Oriens* and apical dendritic tree in *Stratum Radiatum*. In addition to the different neuroanatomical distribution, these trees differ in their molecular composition, e.g., the protein palmitoylation profile^78^, therefore we decided to perform a separate morphometric analysis of dendritic spines located on the basal dendritic tree and apical dendritic tree in the CA1 subregion of the hippocampus. Morphometric analysis revealed that in ANH animals in both basal and apical dendritic trees there was an increase in *length-to-head-width* ratio in comparison to control animals and RES animals (Figure 5D-E, Figure S5). On the other hand, only in the basal dendritic tree in RES animals we observed a trend in the decrease in length-to-head-width ratio in comparison to control animals (p = 0.20) (Figure 5D-E, Figure S5). Moreover, a detailed analysis showed that changes in *length-to-head-width* ratio of RES animals in the basal dendritic tree are associated with a significant increase in the head width and the area of the dendritic spine in comparison to control animals and ANH animals (Figure 5D-E, Figure S5). This result indicates maturation of dendritic spines in this subregion in RES animals, which may be the basis for the mechanism of structural compensation in RES animals. In summary, we conclude that resilient behavior is accompanied by structural compensation in the hippocampus, which is shown in the representative confocal images of dendritic spines in Figure 5E.

**Figure 5.**
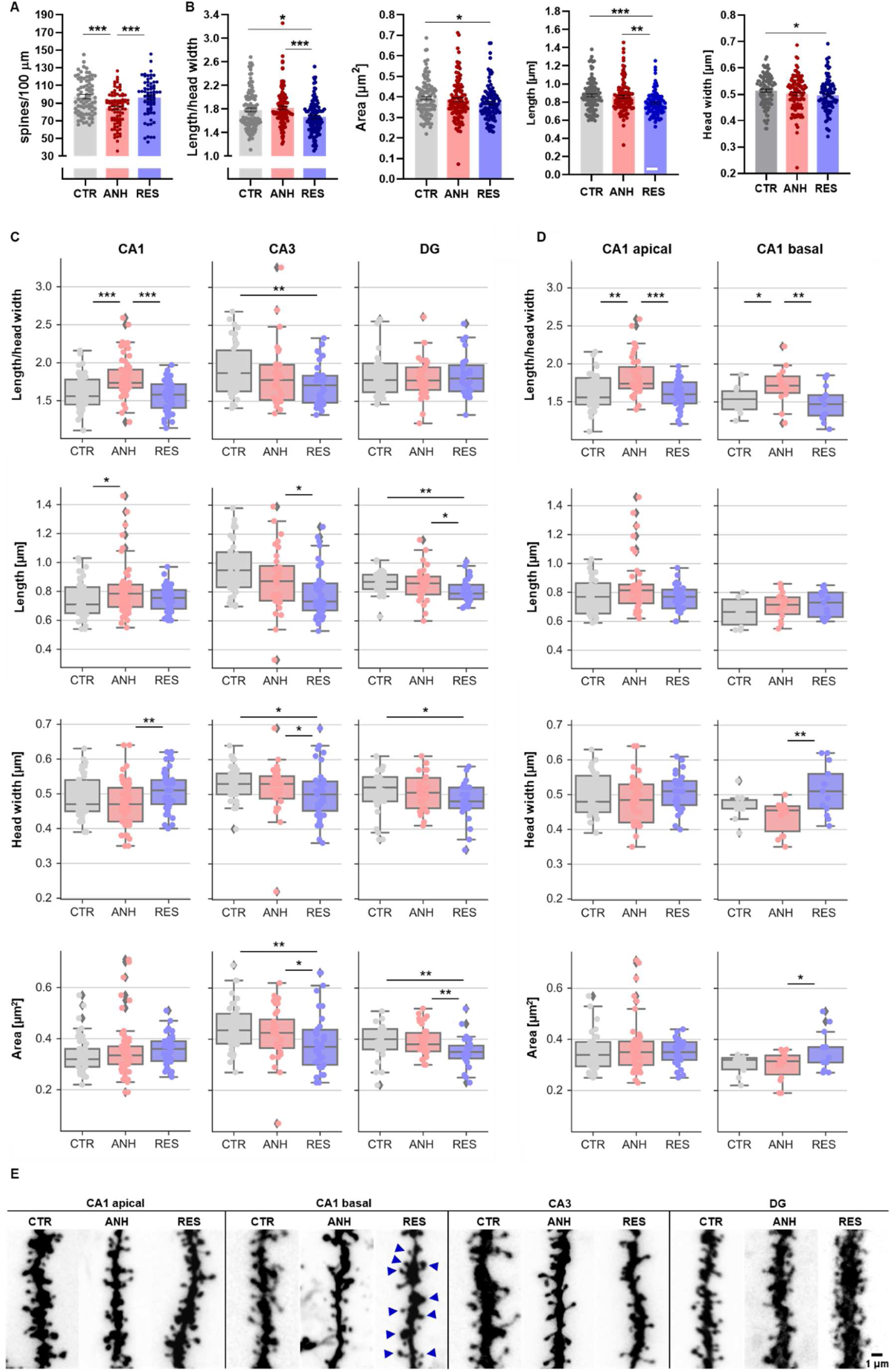
Chronic stress differentially affects the density and morphology of dendritic spines in the hippocampus of CTR, ANH, RES animals, including **(A)** spine density, **(B)** morphology in scale-free parameter (length/head width ratio) of relative morphometric changes, dendritic spine area, length, head width. CTR N_spines_ = 7735, N_dendrites_= 116, N_animals_ = 5; ANH N_spines_ = 7918, N_dendrites_ = 120 N_animals_ = 5; RES N_spines_ = 7026 N_dendrites_ = 107, N_animals_ = 5, **(C)** Morphometric analysis of dendritic spines in the CA1, CA3, and DG subregions of the hippocampi of CTR, ANH, RES animals **(D)** Morphometric analysis of dendritic spines localized in the apical and basal dendrites in the CA1 subregion of the hippocampi of CTR, ANH, RES animals **(E)** representative confocal images of CTR, ANH, RES hippocampal dendritic segments. The arrows indicate the mushroom-shaped dendritic spines in the RES group. Scale bar = 1 µm. Data are presented as the mean ± SEM. The points on the graphs represent the dendrite fragments in 1 field of view; *p < 0.05; **p < 0.01, ***p < 0.001 (nested ANOVA).

## DISCUSSION

In the present study, we aimed to investigate whether stress resilience is an actively-developed process in adult mice. Therefore, we determined the molecular fingerprint of stress-resilient behavior involved in hippocampal neuronal circuits upon chronic stress. We demonstrated that stress resilience is associated with alterations in glutamate receptor signaling pathways, which are exclusively limited to the postsynaptic parts of synapses. We showed that stress resilience manifests itself by compensatory remodeling of dendritic spines, combined with concomitant changes in the S-PALM of synaptic proteins involved in spine morphogenesis, receptor trafficking and glutamatergic neurotransmission.

The complex characterization of synaptic plasticity in resilient animals is an appealing issue. Most of the research on stress resilience is related to a behavioral evaluation in early-life maternal separation models or to the genetic knockouts of targeted proteins that promote stress-resilient behavior in adulthood^5,79,80^. Therefore, we discuss our results with respect to animal models that share similar components of stress procedures to those based on the chronic unpredictable stress and social defeat paradigms. As a result of the multifaceted and comprehensive analysis of the characteristics of the resilient and anhedonic phenotypes, which differ from that of the non-stressed mice, we observed specific changes in synaptic plasticity within behavioral groups. Similar to reports in the literature^81–84^, we observed significant differences in the expression levels of synaptic proteins following chronic stress, but those changes did not differentiate the resilient and anhedonic phenotypes. Importantly, the ClueGO analysis revealed that significantly altered expression levels of the synaptic proteins associated with chronic stress was particularly associated with morphogenesis of dendritic spines and processes related to protein localization to the postsynaptic membrane, which is in agreement with the results of the aforementioned studies.

Several authors noticed robust proteomic alterations in the hippocampus following chronic stress^81,85,86^. However, our results revealed more subtle changes than those reported earlier. This difference may be because we used the synaptoneurosomal fractions and not brain homogenates from the entire hippocampus.

Chronic stress has been shown to affect hippocampal glutamatergic neurotransmission as well as learning and memory^87–89^. In our study, we focused on the general characteristics of the AMPAR- and NMDAR-mediated fEPSPs in each behavioral group. We did not aim to assess how the chronic stress affects LTP and learning and memory in mice. Nevertheless, our CUS protocol was based on that in a study by Strekalova et al.^90,91^ in which behavioral evaluation for learning and memory was performed, showing cognitive impairments in anhedonic animals. This experimental direction is particularly worth investigating in the future, as studies in humans have shown that resilient individuals exhibit cognitive enhancement^92,93^.

The potential cognitive enhancement in the resilient animals may arise from the structural compensation of dendritic spines in the hippocampus (manifested by the higher spine density and levels of the observed spine maturation of already existing spines than those in the anhedonic group). The importance of appropriate restoration of the density and morphology of dendritic spines is emphasized in strategies of MDD treatment^55,57,59,94^. In particular, the results obtained in preclinical studies concerning the activity of fast and long-acting antidepressants, such as glutamate-based antidepressants, e.g., ketamine, or serotonin-based compounds, such as psilocybin, have shown that sustained antidepressant-like effects could causally depend on changes in the density and/or morphology of dendritic spines^95–97^. In general, not every type of stress protocol affects the structure of dendritic spines similarly. In fact, only severe and prolonged exposure leads to pathological remodeling of spines (density and/or morphology), which is assumed to be a structural correlate of depressive symptoms in rodents^98^. In the present study, we have shown that anhedonic animals exhibited a decrease in spine density with no morphometric changes within the whole hippocampus. However, as in the previous study, we have demonstrated that anhedonic behavior is associated with spine elongation exclusively in the CA1 subregion of the hippocampus and that this subregion of the hippocampus is involved in the development of anhedonic behavior^68^. The results obtained in the current study suggest that the CA1 subregion of the hippocampus is also affected by structural and molecular changes during resiliency. Going even deeper, our results suggest that chronic stress differentially affects the structural plasticity of the proximal (basal dendrites localized in in *Stratum Oriens*) and distal (apical dendrites localized in in *Stratum Radiatum*) areas of the hippocampal CA1 subregion. Interestingly, the proximal parts of the dendrites have been shown to have greater regenerative capacity compared to the distal dendrites^99,100^, which is consistent with the observed different structural effects in the basal and apical dendritic trees of pyramidal cells after exposure to CUS. In addition, it is known that, apart from different neuroanatomical distribution, these trees differ in molecular and functional terms^78^. In line with the results of other studies^101–103^, we did not observe differences in the expression levels of synaptic proteins belonging to the mTORC1 complex following chronic stress. Nevertheless, the reduced level of Rictor protein (mTORC2 binding partner) in the resilient group gained our attention due to mTORC2 insensitivity to the rapamycin^104^. The role of mTORC2 in the hippocampal synaptic plasticity has been widely reported^105–107^. However, how its complex may be engaged in the behavioral stress response and how it is palmitoylated have not yet been studied. Notwithstanding, we also observed differences in the S-PALM (Cys-162) of tetratricopeptide repeat ankyrin repeat and coiled-coil containing 2 (TANC2) between the resilient and anhedonic animals and the controls. Deletion of TANC2 in the hippocampus hyperactivates mTORC1/mTORC2-dependent signaling pathways, leading to cognitive impairment and hyperactivity in mice^108^. Moreover, TANC2 directly interacts with postsynaptic density protein 95 (PSD-95) and constitutes an endogenous inhibitor of mTORC1/mTORC2 complexes^108^. Moreover, it has been shown that ketamine activates the mTORC1 complex by suppressing the interaction of TANC1/2 with mTOR interaction but does not affect the interaction of TANC1/2 with PSD-95^108^. Despite the still unknown physiological role of S-PALM of TANC2 (Cys-162), its role in the biological basis of stress resilience should be considered as a potential molecular target in future studies.

In addition to the involvement of the mTOR complex in the regulation of dendritic spine structure, as well as glutamatergic neurotransmission, GSK-3-beta kinase also plays a decisive role in the development of depressive-like behavior, and the structural remodeling of dendritic spines^109–111^. In particular, our results have revealed the possible role of GSK-3-beta kinase in the genesis of resilient behavior. We observed a decreased level of beta-catenin (responsible for cell survival in a Wnt-dependent manner), suggesting the impact of GSK-3-beta/APC complex activity in the resilient group. The distinct S-PALM of two cysteines (Cys-912 and Cys-2664) in adenomatous polyposis coli (APC) protein was also observed in resilient and anhedonic mice. Due to the formation of a complex of APC and GSK-3-beta kinase, leading to the degradation of beta-catenin in the proteasome^112^, the S-PALM of APC might be involved in the regulation of beta-catenin levels. Moreover, we observed a decrease in the level of the aforementioned Rictor protein in the resilient group, which was negatively correlated with the functioning of GSK-3-beta kinase^113^. Thus, we can speculate that enhanced S-PALM of APC can negatively regulate the formation of the APC/GSK-3-beta complex, decreasing the degradation of proteins, such as beta-catenin or Rictor, in the anhedonic phenotype. However interesting, these outcomes should be interpreted carefully. First, the level of beta-catenin is not regulated only by its degradation in the proteasome; therefore, the correlation between the expression of beta-catenin and the activation of the GSK-3-beta/APC complex could be misleading^112^. Concurrently, GSK-3-beta kinase activity is much more complicated than its simple involvement in the complex with APC^112^. In particular, the interplay between GSK-3-beta kinase and mTOR complexes, such as TSC-1/TSC-2, is not dependent on the GSK3-beta/APC complex but on the dephosphorylation of GSK-3-beta (Ser-9)^112,114^. Therefore, properly understanding the role of GSK-3-beta kinase in resilient and anhedonic mice requires more profound studies with complex profiling of the phosphorylation of synaptic proteins. Despite the unclear role of GSK-3-beta kinase in the decreased level of beta-catenin in the resilient phenotype, the role of this protein in the development of stress resilience is still interesting. Vidal et al. showed that inhibition of beta-catenin in GLAST-expressing cells lead to the development of depressive-like behavior while its stabilization led to a resilient state in a model of chronic exposure to corticosterone^115^. Therefore, the decreased level of beta-catenin in the hippocampus of resilient animals should be considered in the context of other signaling pathways because, as we showed, different cellular events underlie stress resilience and depressive-like behavior. The NMDAR and AMPA receptor subunits revealed alterations in S-PALM following chronic stress. In particular, we observed increased levels of the palmitoylation of the GluA1 subunit of AMPA receptors at two cysteines (Cys-601 and Cys844). As previously reported, palmitoylation of these cysteine residues increases the anchoring of the receptor GluA1 subunit to the Golgi apparatus and inhibits the interaction between the subunit and the synaptic 41N protein, affecting glutamatergic transmission^116,117^. Increased palmitoylation of the Cys-202 residue of the GDP dissociation inhibitor-1 (GDI-1) exclusively in the anhedonic group may also indirectly regulate the turnover of AMPA receptors anchored to dendritic spines and may explain the observed decrease in spine density. It was shown previously^118,119^ that the interactions of GDI-1 with the Rab family proteins are responsible for maintaining equilibrium between exocytosis and endocytosis through LTP. Unfortunately, the physiological role of Cys-202 palmitoylation of GDI-1 remains unknown. We also observed an increased level of palmitoylation of two cysteines (Cys-954 and Cys-1173) in the cytoplasmic domain of the GluN2B subunit in the anhedonic group. However, the physiological role of these GluN2B modifications has also not been described. Nevertheless, the differences in the functional readout of AMPA and NMDA receptors upon chronic stress could also be explained by the fact that palmitoylation is not the only posttranslational modification that occurs at the synapse. Several serine residues in the GluA1-A4 subunits of the AMPA receptor, as well as the GluN2B subunit, undergo phosphorylation, affecting receptor trafficking, conductance, and the frequency of the channel opening^120–122^. Therefore, the interplay between the expression, palmitoylation, phosphorylation, and ubiquitination of synaptic proteins produces a functional effect.

The mechanism underlying region-specific differences in the function of excitatory synaptic transmission in CA1 and CA3 regions in anhedonic and resilient animals remain elusive (Fig. 4, Fig. S3, Fig. S4). At least in CA1 region, most likely the postsynaptic and not presynaptic site is the locus for molecular fingerprint of resilience. It is possible, that the palmitoylation of AMPA and NMDA receptor subunits along with GDI-1 might be differentially affected in resilient and anhedonic animals. Whether and how these processes are manifested within behavioral groups is a matter of further study.

In conclusion, we demonstrated that stress-resilient behavior in adult animals is a actively regulated process accompanied by a set of unique functional, proteomic and morphological features in the brain. In particular, we have shown that the most robust synaptic alterations underlying stress resilience are associated with the structure of dendritic spines and postsynaptic intracellular signaling pathways in the hippocampus. At the cellular level, these differences might be triggered by S-PALM of synaptic proteins and translated into the regulation of synaptic receptors. However, further studies are required to indicate the chemical kinetics of these processes and their role in the behavioral stress response.

## MATERIALS AND METHODS

### Animals

Ten-week-old male C57BL/6J mice (Medical University of Bialystok, Poland) were individually housed under a reverse 12/12 h light/dark cycle (lights on at 8:00 PM) with food and water available *ad libitum*. Male 12-week-old CD1 mice (Janvier Labs, France) were used as resident intruders in the social defeat stress procedure and kept with the stressed C57BL6J mice in the same animal room. Male 12-week-old Wistar rats (Mossakowski Medical Research Institute, Polish Academy of Sciences, Warsaw, Poland) were used for predator stress. Male 12-week-old C57BL/6J mice (Mossakowski Medical Research Institute, Polish Academy of Sciences, Warsaw, Poland) were used for isolation of CA1 *Stratum Radiatum* and *Stratum Oriens* subregions of hippocampus. All animal procedures were performed according to the guidelines of the Polish Ethical Committee on Animal Research (permission no. 132/2016, 2011/2020, 203/2021, 204/2021).

### Mouse Model of Stress Resilience based on Chronic Unpredictable Stress (CUS)

To evaluate depressive-like behavior in the mouse model, we used the chronic unpredictable stress paradigm (CUS) and behavioral evaluation as we described previously^29,68^. We have described a highly reproducible protocol with all the relevant technical details previously ^69^. Briefly, before CUS, C57BL/6J mice were subjected to 2 weeks of room acclimatization, consisting of 1 week of handling. Mice were weighed and their baseline sucrose preference (SPT0) was measured before the CUS procedure. Then, based on their baseline parameters, mice were assigned to a control and stress group housed in two separate rooms. The 2-week CUS protocol consisted of 2 out of 3 different types of stressors chosen in a semirandom manner and applied daily during the dark phase under red light in the following sequence of restraint stress, tail suspension, and social defeat stress, with an intersession of at least 3 h. During each light phase during the stress protocol, the mice were exposed to a rat. To stabilize glucocorticoid levels after the last exposure to a stressor, the mice were left undisturbed overnight before beginning the sucrose preference test. Thus, 16 h after the last stressor, the mice underwent a sucrose preference test (SPT1), resulting in the determination of sucrose preference 24 h after the last stressor, and thereafter, the body weight measurements and the forced swim test were performed. All mice were sacrificed 12-16 h after the behavioral evaluation (36-38 h after the last stressor). To correlate the molecular, functional, and structural readouts of excitatory synaptic plasticity in the hippocampus in relation to animal behavior, two independent CUS experiments were performed in which within each animal, one hippocampus was subjected to synaptoneurosome isolation (mass spectrometry analysis) or DiI labeling (dendritic spine imaging), and the second hippocampus was subjected to electrophysiology. The experimental design is outlined in Figure 1A.

#### Restraint stress

The mice were placed inside a plastic tube (26 mm internal diameter) for 2 h during the dark phase.

#### Tail suspension stress

The mice were subjected to the tail suspension procedure by being hanged from the tails with adhesive tape for 40 min during the dark phase. To prevent the mice from climbing their tails, plastic cylinders (4 cm x 0.5 cm) were placed at the base of their tails.

#### Social defeat stress

During each 30 min social defeat session performed in the dark phase, aggressive CD1 animals were placed in the home cages of C57BL/6J mice in the stress group. CD1 aggressive mice were selected as the CD1 mice that attacked C57BL/6J mice in less than 60 s without injuring them. During each session, the C57BL/6J mice exhibited signs of social defeat stress, such as a flight response, submissive posture, and audible vocalization. If the mice in the stress group did not display signs of social defeat stress, then the CD1 mouse was changed to another CD1 mouse. In rare cases of physical harm that occurred between pairs of mice, aggressive CD1 individuals were immediately removed from the cage of the C57BL/6J resident mice.

#### Predator stress

The mice were individually introduced into transparent, well-ventilated cylinders (15 cm × 8 cm) with food and bedding. The cylinders were then placed for 12 h (08:00 P.M. – 08:00 A.M.) into a rat home cage that contained a rat during the light phase. For the rest of the day (08:00 A.M. – 08:00 P.M.), the mice and rats were housed in their home cages in the same experimental room.

## Behavioral Tests

### Sucrose preference test (SPT)

Mice were given free-choice access to 1% sucrose solution and water that were provided in identical bottles for 8 h during the dark phase under a reverse light dark cycle. The percentage of sucrose preference was calculated as follows:

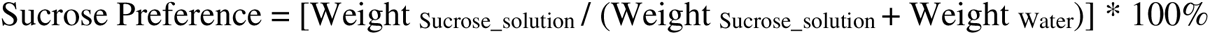

The consumption of water and sucrose solution was estimated simultaneously in the control and experimental groups by weighing the bottles. To eliminate possible bias from side preference, the positions of the bottles were changed after 4 h of the test. Twenty-four hours before the baseline sucrose preference test performed before the CUS procedure (SPT0), 2.5% sucrose solution was given to all animals for 2 h to prevent the possible effects of taste neophobia. The other conditions of the test were performed as previously described^29,68,90,91^. Sucrose preference after CUS (SPT1) values of < 70.7% in mice in the stress group, measured 24 h after the last stressor, was the criterion for “anhedonia”, defined by the difference between the control and stressed groups > 2 × SD. Anhedonic mice were previously shown to display depressive-like behavior^90,91^. None of the control animals exhibited < 70.7% sucrose preference in SPT1. Stressed mice with sucrose preference > 70.7% at the end of the CUS experiment were defined as resilient animals. The SPT evaluation before (SPT0) and after CUS (SPT1) is outlined in Figure 1B.

### Forced swim test (FST)

Each mouse was placed into a cylindrical glass containers (20 cm × 40 cm) filled with warm water (∼27 °C) for a 6 min swimming session. The test was conducted under red light during the dark phase of the light/dark cycle after 1 h of room acclimatization where the behavioral test was performed. The sum of the floating time during the last 4 min was measured by visual scoring offline and defined as the time spent immobile.

All behavioral scorings were performed blind to the treatment groups.

### Acute Brain Slice Electrophysiology

Acute hippocampal brain slices were obtained from control, anhedonic, and resilient mice (N at least 5 mice/group) according to the protocol described previously^76^. The hippocampi from one hemisphere were dissected and cut into 350 µm thick slices using a vibratome (VT1200S, Leica, Germany) in ice-cold buffer that contained 75 mM sucrose, 87 mM NaCl, 2.5 mM KCl, 1.25 mM NaH_2_PO_4_, 25 mM NaHCO_3_, 0.5 mM CaCl_2_, 10 mM MgSO_4_*7H_2_O, and 20 mM glucose, pH 7.4. Slices were recovered in the same solution for 15 min (32 °C) and were subsequently stored in oxygenated (95% O_2_, 5% CO_2_) artificial cerebrospinal fluid (aCSF) that contained 125 mM NaCl, 25 mM NaHCO_3_, 2.6 mM KCl, 1.25 mM NaH_2_PO_4_, 2.0 mM CaCl_2_, and 20 mM glucose, pH 7.4. Recordings were made in aCSF after 2 h of slice recovery. Schaffer collateral pathway or commissural/associational-CA3 synapses were stimulated with a concentric bipolar electrode at CA1 or CA3 subregions, respectively (0.1 Hz, 0.3 ms, FHC, Bowdoin, ME USA). Compound AMPAR- and NMDAR-mediated fEPSPs were recorded with glass micropipettes that were filled with aCSF (1-3 M Ω resistance) in the *Stratum Radiatum* of the CA1 or CA3 subregion of the hippocampus as described previously^88,134^. NMDAR-mediated signals were isolated from compound fEPSPs with the AMPA/kainate receptor antagonist DNQX (20 μM) and L-type calcium channel blocker nifedipine (20 μM) in Mg^2+^-free solutions, as described previously^76^. At the end of each recording, the NMDAR antagonist APV (50 μM) was used to confirm the origin of the recorded fEPSPs. All of the drugs were obtained from Sigma-Aldrich (Poland) and Tocris (UK). Input–output (I–O) relationships were built for fEPSPs amplitudes upon monotonically increasing the stimuli in the range of 0–300 µA (13 points, applied once at 0.1 Hz). Baseline stimulation was set at 0.1 Hz, and for baseline and paired-pulse stimulation protocols (interstimulus interval 25 ms), the stimulation strength was set to 40% of the maximum fEPSP amplitude. For better data visualization, the input–output curves shown in Figure 4A were fitted with Equation:

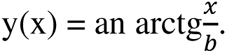

Once the data were fitted, more complicated dependences, e.g., fEPSP amplitude vs. fiber volley amplitude, were recovered by combining the fit in the form of z(x) (with parameters a, b) with an inverse of the relation (1) in the form of x(y), with 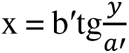, which led to 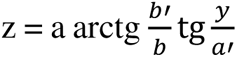(Figure 4B-C).

Due to the fact, that the data do not fulfill the assumptions of mixed design ANOVA test, and because of the complex design of the experiment, we obtained the p-value using a randomization approach by Monte Carlo methods^123^. We computed statistical significance of the differences (two-tailed tests) between the stimulation current, fiber volley amplitudes and fEPSPs, *V(I)*, obtained from the electrophysiological experiments (Figure 4A-C). The p-values were calculated as follows: In the first step, we quantified the differences between the measured *V^A^(I_i_)* and *V^B^(I_i_)* curves using the *L_2_* norm, defined as

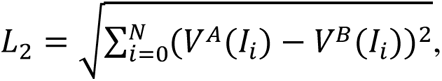

where *I_i_* is the stimulation current for the *i-*th measurement point and *N* is the total number of the measurement points. To compute the p values, we created the null-hypothesis ensemble, using the subject randomization and bootstrap techniques^124^. The p value for the difference between two *V(I)* curves was calculated as

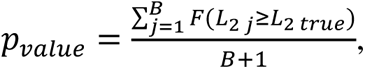

where *F* is the indicator function that takes the value one when its argument is true and zero when it is false, *L_2 j_* is the *L_2_* norm for the *j-*th element of the null-hypothesis ensemble, *L_2 true_* is the actual value of *L_2_* for the analyzed *V(I)* curves, and *B* is the number of randomizations (we used *B* = 1000).

The electrophysiology data were analyzed using AxoGraphX software as described previously^76^.

### Synaptoneurosomes

After euthanasia by cervical dislocation, the mice were decapitated, and hippocampi were removed and homogenized with a Dounce homogenizer in 3 mL of buffer A (5 mM HEPES (pH 7.4), 0.32 M sucrose, 0.2 mM ethylenediaminetetraacetic acid (EDTA), 50 mM N-ethylmaleimide (NEM), and protease inhibitor cocktail. Nuclei and cell debris were pelleted by 5 min centrifugation at 2 500 × g. The supernatant was then centrifuged at 12000 × g for 5 min. The obtained pellet fraction was layered over a discontinuous Ficoll (Sigma-Aldrich) gradient (4%, 6%, and 13%) and centrifuged at 70000 × g for 45 min. The synaptoneurosomal fraction was collected in buffer A and centrifuged at 20000 × g for 20 min. The pellet corresponded to the synaptoneurosomes fraction. Purified synaptoneurosomes were obtained from the hippocampus collected from one hemisphere of control, anhedonic, resilient animals (N mice/group = 5) and subjected to mass spectrometry analysis as described previously^72^. The obtained synaptoneurosomal fraction was visualized using electron microscopy and is presented in the Figure S2.

### Mass Spectrometry

The S-PALM or total protein peptide mixture (20 µL) was applied to the nanoACQUITY UPLC Trapping Column (Waters, 186003514) using water containing 0.1% formic acid as the mobile phase and transferred to the nanoACQUITY UPLC BEH C18 Column (75 µm inner diameter; 250 mm long, Waters, 186003545) using an acetonitrile gradient in the presence of 0.1% formic acid with a flow rate of 250 nL/min. The column outlet was directly coupled to the ion source of the Thermo Orbitrap Elite mass spectrometer (Thermo Electron Corp., San Jose, CA, USA) working in the regime of data-dependent MS to MS/MS switch. HCD fragmentation was used. All MS runs were separated by blank runs to reduce the carry-over of peptides from previous samples. The results of measurements were processed using Mascot-Distiller 2.7.1 software (MatrixScience, London, UK, on-site license). The Mascot search engine (version 2.7.1) was used to compare data against the UniProtKB/Swiss-Prot database (Swissprot 2020_02; 16,905 sequences). The search parameters were set to the following: taxonomy (*Mus musculus*), variable modifications – cysteine carbamidomethylation or N-malemideidation, methionine oxidation, peptide tolerance (5 ppm), fragment mass tolerance (5 ppm). Enzyme specificity was set for trypsin with one missed or nonspecific cleavage permitted. The mass calibration and data filtering described above were also carried out. The lists of the peptide sequences (SPL) that were identified in all of the LC–MS/MS runs from synaptoneurosomal fractions were merged into one peptide list using MascotScan software (http://proteom.ibb.waw.pl/mscan/, accessed on 9 April 2021). The SPL consists of sequences of peptides with Mascot scores exceeding the threshold value corresponding to a 5% expectation value and FDR of 1% calculated by the Mascot procedure. For proteome quantitative analysis, peptide intensities were determined as the surface of the isotopic envelope of the tagged isotopic envelopes. Before the analysis, quantitative values were normalized with LOWESS as described previously^72^.

### Functional Bioinformatics Analysis of Mass Spectrometry Data

For integrative analysis, we used ClueGO software to observe differential proteins involved in the GO terms. The input list of proteins for each GO analysis was distinguished on the basis of proteomic data analysis and Venn diagram analysis. The lists of proteins are grouped in Supplementary Tables 1-2 included in the Supplementary Materials. Proteins were analyzed with ClueGO v2.6.4/CluePedia v1.6.5 to obtain complete Gene Ontological terms (GO) from our datasets. ClueGO integrates GO terms and creates an organized GO/pathway term network. The statistical test used for the node enrichment was based on a right-sided hypergeometric option with a Benjamini–Hochberg correction and kappa score of 0.5. As a reference set for term enrichment calculations, we utilized genes from the *Mus musculus* genome (NCBI unique Gene identifiers). GO enrichment was conducted for different sets of proteins, and p values < 0.05 were considered to be significant. ClueGO results are grouped in Figure 5.

### DiI Staining of Brain Slices and Morphometric Analysis of Dendritic Spines

To visualize changes in the structure of dendritic spines, 1,10-dioctadecyl-3,3,3,30-tetramethylindocarbocyanine perchlorate (DiI) staining was performed on one brain hemisphere fixed by incubation for 1 h in 1.5% paraformaldehyde. The hemispheres were sliced using a Leica vibratome. Slices (140 µm thick) containing hippocampal structures were allowed to recover for at least 1.5 h at room temperature. Random cell labeling was performed using 1.6 µm tungsten particles (Bio–Rad, Hercules, CA, USA) that were coated with a propelled lipophilic fluorescent dye (DiI; Invitrogen) delivered to the cells by gene gun (Bio–Rad) bombardment. Images of hippocampal neurons covered with dendritic spines were acquired under 561 nm fluorescent illumination using a confocal microscope Zeiss LSM800 (63× objective, 1.4 NA) at a pixel resolution of 1024 × 1024 with a 2.4 × zoom, resulting in a 0.07 µm pixel size. The analysis of dendritic spine structure and calculation of changes in spine parameters were performed as described previously^29,68,125^. The images that were acquired from the brain slices were processed using ImageJ software (National Institutes of Health, Bethesda, MD, USA) and analyzed semiautomatically using custom-written SpineMagick software (patent no. WO/2013/021001). The analyzed dendritic spines belonged to secondary and ternary dendrites to reduce possible differences in spine morphology caused by the location of spines on dendrites with different ranks. To quantify the changes in spine shape, we analyzed the relative changes in the spine length-to-head-width ratio (the scale-free parameter). The spine length was determined by measuring the curvilinear length along a fitted virtual skeleton of the spine. The fitting procedure was performed by looking for a curve along which integrated fluorescence was at a maximum. Head width was defined as the diameter of the largest spine section while excluding the bottom part of the spine (1/3 of the spine length adjacent to the dendrite). Dendritic segments of 5 animals per group (70-93 cells/group) were morphologically analyzed resulting in CTR N*_spines_* = 7 735, N*_dendrites_* = 116; ANH N*_spines_* = 7918, N*_dendrites_* = 120; RES N*_spines_* = 7026 N*_dendrites_* = 107. To assess dendritic length for spine density analysis, we measured curvilinear length along the analyzed dendritic segment, which was obtained by fitting an n-order polynomial resulting in CTR*_dendritic length_* = 23795.67 µm, ANH*_dendritic length_* = 27179.45 µm, and _RES*dendritic length*_ = 21996.42 µm. The spine number was counted manually by a trained neurobiologist in ImageJ software. The statistical analysis was performed using nested analysis of variance (including the number of animals and the number of dendrite fragments in 1 field of view/spines analyzed per behavioral group. The distributions of spine parameters for spine density, length/head width, and area for the control, anhedonic and resilient groups are presented in Figure S4.

### Acyl-Biotin Exchange (ABE)

The acyl-biotin exchange (ABE) method was used to analyze protein palmitoylation in tissue samples from CA1, CA3 and DG subregions of the hippocampus of resilient, anhedonic and control animals. First, the samples were homogenized using Dounce homogenizer in a buffer containing 50 mM Tris HCl (pH 7.5), 150 mM NaCl, 1 mM EDTA, 4% SDS and 1% Triton X-100 (ABE buffer). The extracted proteins were reduced with 10 mM tris(2-carboxyethyl)phosphine (TCEP) and incubated with 50 mM N-ethylmaleimide (NEM) at 4 °C for 16 h to block free thiol groups. Next, to remove the excess of unreacted NEM the proteins were precipitated with 96% ice-cold ethanol. The pellets were resuspended in the ABE buffer and the samples were treated with 400 µM thiol-reactive biotinylation reagent HPDP-biotin (N-[6-(biotinamido)hexyl]-3’-(2’-pyridyldithio)propionamide) and subsequently split into two equal parts. One part was treated with 1 M hydroxylamine to cleave thioester-linked palmitoyl moieties and expose freshly formed-thiols to HPDP-biotin; the other part was treated with Tris buffer pH 7.5 as a control which is used to identify unspecific binding of the HPDP-biotin. The samples were incubated in the dark for 1.5 h with constant agitation at room temperature. Finally, aliquots of the samples were subjected to SDS-PAGE and Western blotting for the analysis of the total palmitoylation (visualization of all biotinylated proteins) Stain-free total protein visualization was used as a loading control for total palmitoylation.

### Western blot analysis

Protein samples were separated by SDS–PAGE and transferred to polyvinylidene difluoride membranes (Immobilon-P, Millipore). The membranes were then blocked with 10% nonfat milk in Tris-buffered saline with 0.1% Tween 20 (TBST). For the visualization of the total palmitoylation (biotinylated proteins after ABE) blocked membranes were incubated for 2 h in room temperature with Horseradish Peroxidase Avidin D (1:4000, A-2014, Vector Laboratories) diluted in 5% nonfat milk in TBST. After washing, the bands were detected using the SuperSignal West Femto Maximum Sensitivity Substrate (for CDC42 and 5-HT7R detection) or SuperSignal West Pico PLUS Chemiluminescent Substrate (for GAPDH detection) (Thermo Fisher Scientific).

### Statistical Analysis

Statistical analysis was performed using GraphPad Prism8 software (GraphPad, San Diego, CA, USA). One-way or two-way analysis of variance (one-way or two-way ANOVA) followed by post hoc tests was used for multiple comparisons to identify significant differences between experimental groups. In the case of unequal variances, the Welch correction was applied. If the data were not normally distributed, the Mann–Whitney test was used. Behavioral data were analyzed using one-way ANOVA followed by the Bonferroni or Tukey post hoc tests. For the statistical analysis of the structural plasticity of dendritic spines (density and morphology), we used nested-ANOVA statistics, including the number of animals and the number of dendrite fragments in 1 field of view /spines analyzed per behavioral group. The nested-ANOVA statistics were performed with R 4.1.1, the language and environment for statistical computing (R Foundation for Statistical Computing, Vienna, Austria). The zymograms were quantitatively analyzed by the sum of replicates (each data point on a replicate is divided by the sum of the values of all data points in that replicate)^126^. A p value of < 0.05 was considered statistically significant for all tests except for mass spectrometry-based palmitoylation analysis results in which < 0.01 was used. The data are presented as the mean value ± SEM. All analyses were performed in a blinded manner. Information about the statistical test used is in the legend for each figure.

### Data availability

All data supporting the findings of this study are available within the article or are available from the corresponding author upon reasonable request. The mass spectrometry proteomics data have been deposited to the ProteomeXchange Consortium via the PRIDE partner repository with the dataset identifier PXD026590.

### Author contributions

Conceived the study: JW, EB and MB. Experimental design: JW, MB, EB, AK, MZK, MM, TW. Performed the experiments: EB, AK, TW, MM, MZK, KB, KHO, ADW, MB, JL. In detail: chronic unpredictable stress model (CUS): EB, AK, KB, KHO, ADW, and SA; behavioral tests: EB, AK, BP; isolation of hippocampal synaptoneurosomes: MZK, IF; mass spectrometry: MZK; electrophysiology: TW; DiI labeling of hippocampal slices: MM, MR, EB; image acquisition: MM, MR; morphometric analysis of dendritic spines: EB; acyl-biotin exchange: AP, KB; analyzed the data: EB, AK, MZK, JW, JM, MB, AP, JL; statistical analysis: BR, RW, PS, EB, MB. Contributed reagents/materials/analysis tools/data interpretation: JW, EB, MB, BR, RKF, BP, BS, KR, JD, PJ, KR, EP. Wrote the paper: EB, JW, BP, MB. All authors corrected and accepted the manuscript.

## Acknowledgments

This work was supported by grants from a National Science Centre (UMO-2021/41/B/NZ4/02603, UMO-2017/26/E/NZ4/00637 and UMO-2019/35/D/NZ4/02042). TW, AP were supported by a National Science Centre (UMO-2019/34/E/NZ4/00387). KB was supported by a National Science Centre (UMO-2020/37/N/NZ4/02869). AD was supported by a National Science Centre grant (UMO-2018/29/B/NZ4/01473). JL was supported by grant from Deutsche Forschungsgemeinschaft (LA4465). EP was supported by grant from Deutsche Forschungsgemeinschaft (PO732).

## Declaration of Interests

The authors declare that they have no conflicts of interest.

## SUPPLEMENTAL FIGURES

**Figure S1.**
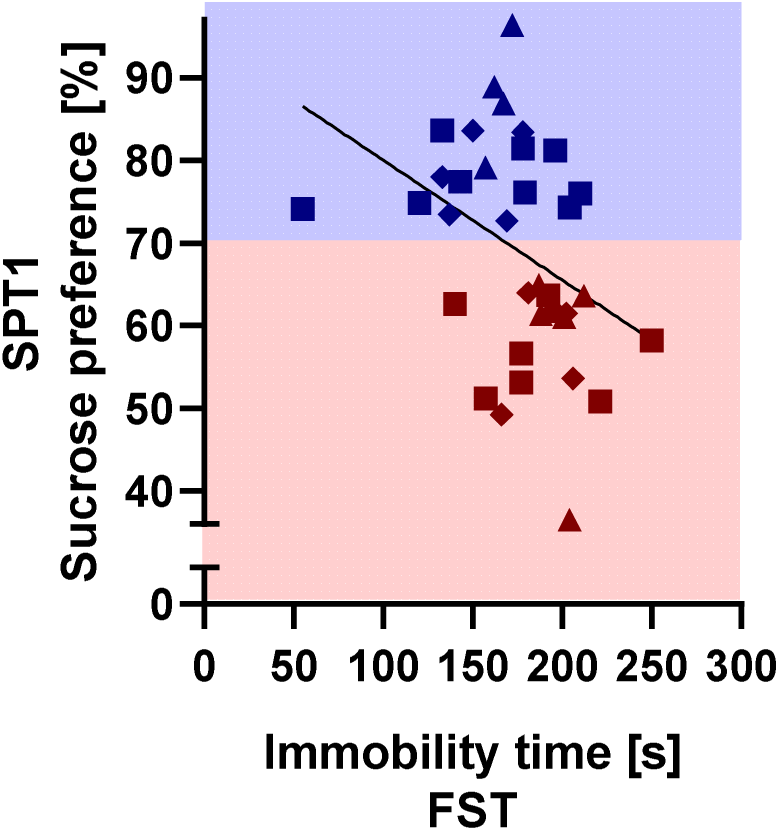
Correlation in the stress group between the behavioural outcomes in sucrose preference test (SPT1) and forced swimming test (FST). Pearson’s correlation coefficient=-0.3771, R squared=0.1422; p=0.02; n = 37.

**Figure S2.**
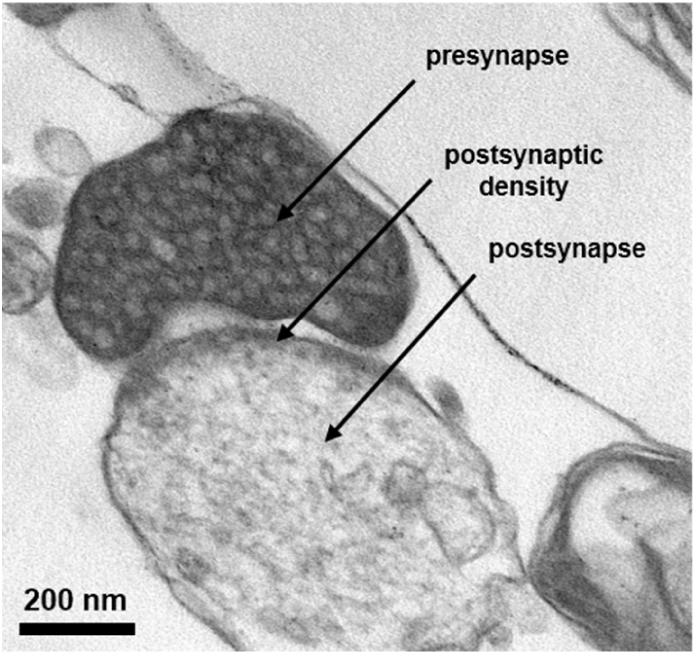
Identification of presynaptic and postsynaptic parts in the synaptoneurosomal hippocampal fraction using electron microscopy.

**Figure S3.**
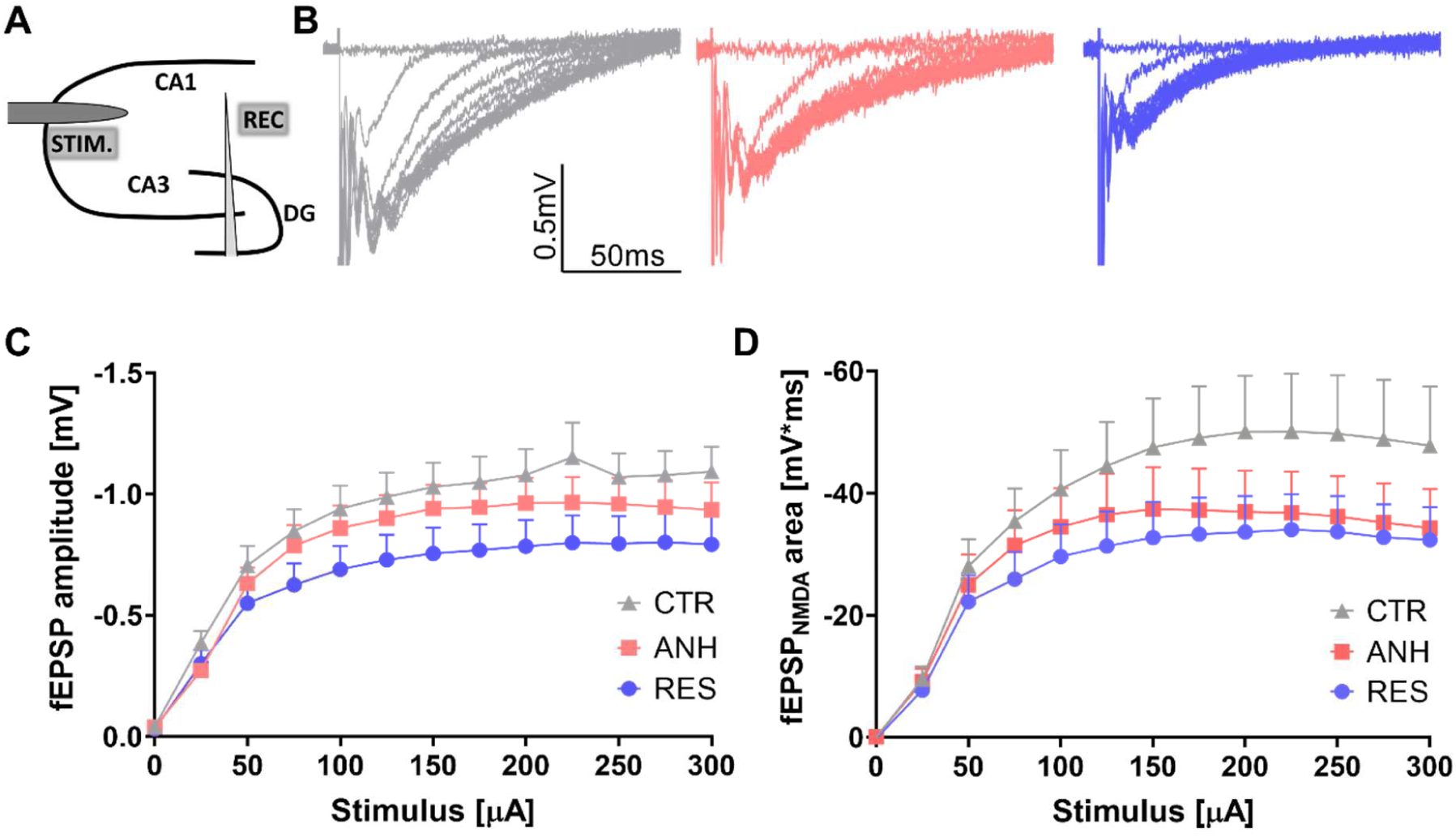
**(A)** Scheme of electrophysiological recordings **(B)** Exemplary compound fEPSPs evoked in the CA1 region of acute hippocampal slices in response to monotonically increased current stimuli in control (gray), anhedonic (red) and resilient group (blue). **(C-D)** Average input–output relationships for AMPAR-mediated **(C)** and NMDAR-mediated **(D)** fEPSPs. The average fEPSP amplitude and area did not differ between groups (CTR vs RES p=0.9, CTR vs ANH p=0.08, ANH vs RES p=0.16 for fEPSP amplitudes, and CTR vs RES p=0.8, CTR vs ANH p=0.37, ANH vs RES p=0.55 for fEPSP area, respectively, Monte Carlo simulations, n_slices_=15-20 (C) and n_slices_=12-16 (D), N_animals_=5-6 per group, for all comparisons). Data are presented as the mean ± SEM.

**Figure S4.**
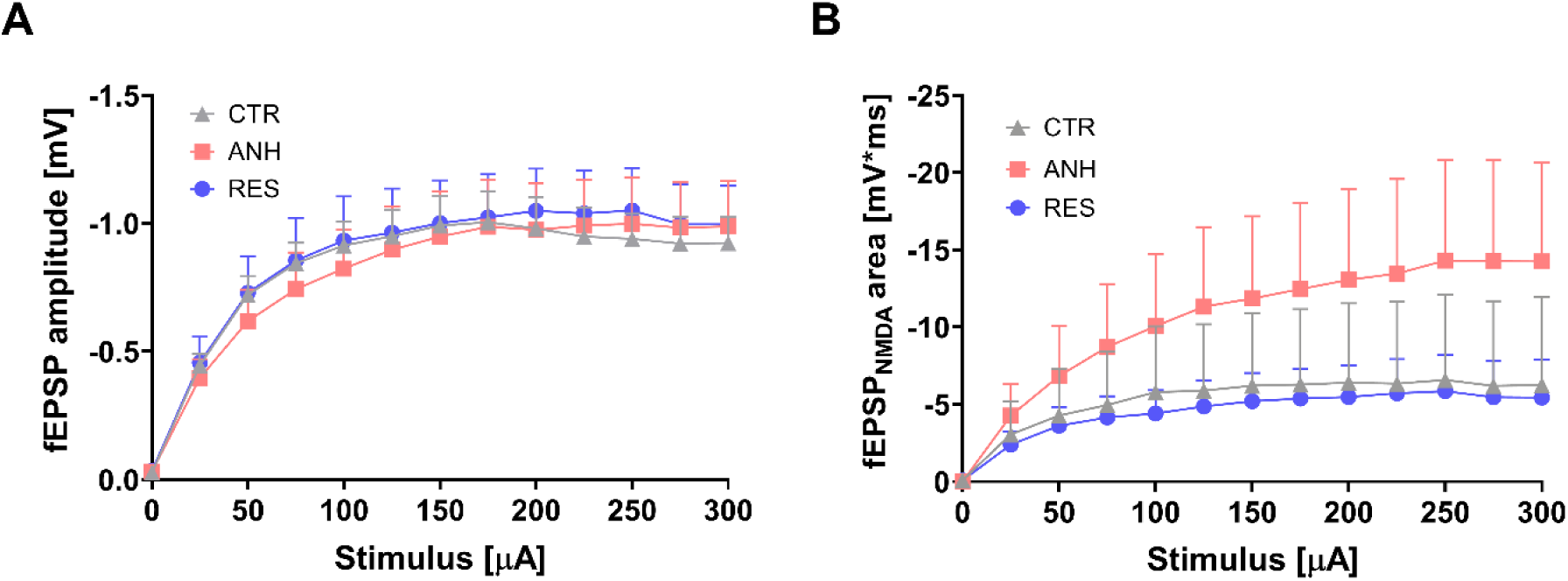
Chronic stress differentially affects NMDAR-mediated neurotransmission in the CA3 region of the hippocampus. **(A-B)** Average input–output relationships for AMPAR-mediated **(A)** and NMDAR-mediated **(B)** fEPSPs evoked in the CA3 region of acute hippocampal slices in response to monotonically increased current stimuli. **(A)** CUS did no results in any change in fEPSP amplitudes (n_slices_=6-8, N_animals_=4-6 per group, p>0.7 for all comparisons, Monte-Carlo simulations). **(B)** The area of the NMDAR-mediated component of fEPSPs was significantly upregulated in Anhedonic compared to Resilient group (p=0.031) while other groups did not differ significantly (n_slices_=3-5, N_animals_=3-6 per group, CTR vs RES p=0.87, CTR vs ANH p=0.085, Monte-Carlo simulations). Data are presented as the mean ± SEM.

**Figure S5.**
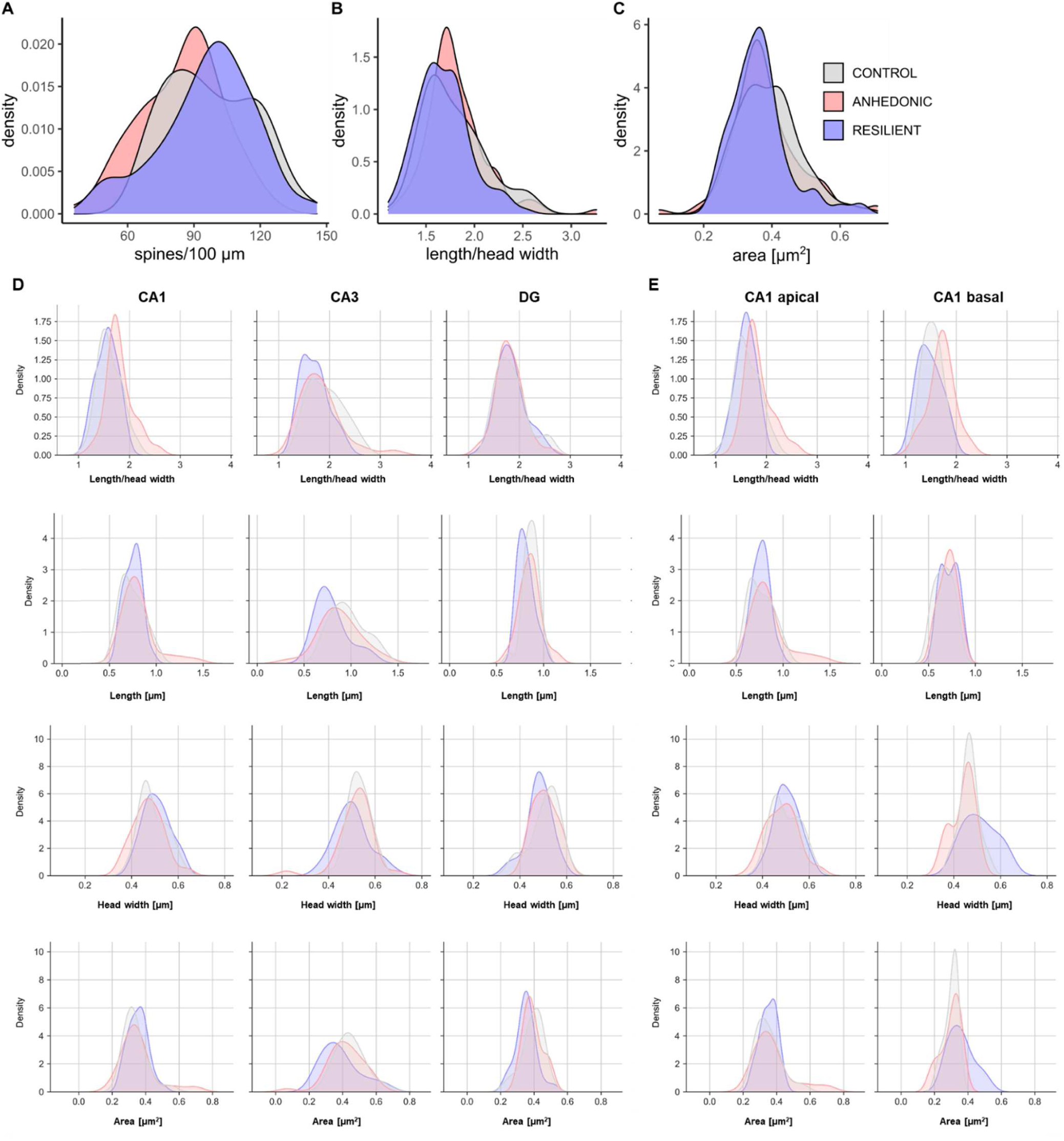
Distributions of (A) spine density, (B) length/head width, and (C) area of dendritic spines in the hippocampus of CTR, ANH, RES animals. (D) Distributions of length/head width, length, head width and area of dendritic spines in the CA1, CA3 and DG subregions of hippocampus of CTR, ANH, RES animals. **(E)** Distributions of length/head width, length, head width and area of dendritic spines localized in the apical and basal dendrites in the CA1 subregion of the hippocampi of CTR, ANH, RES animals. The distributions show that the density of probability integrated with the variable on the x-axis is normalized to unity.

**Supplemental Table 1A.**
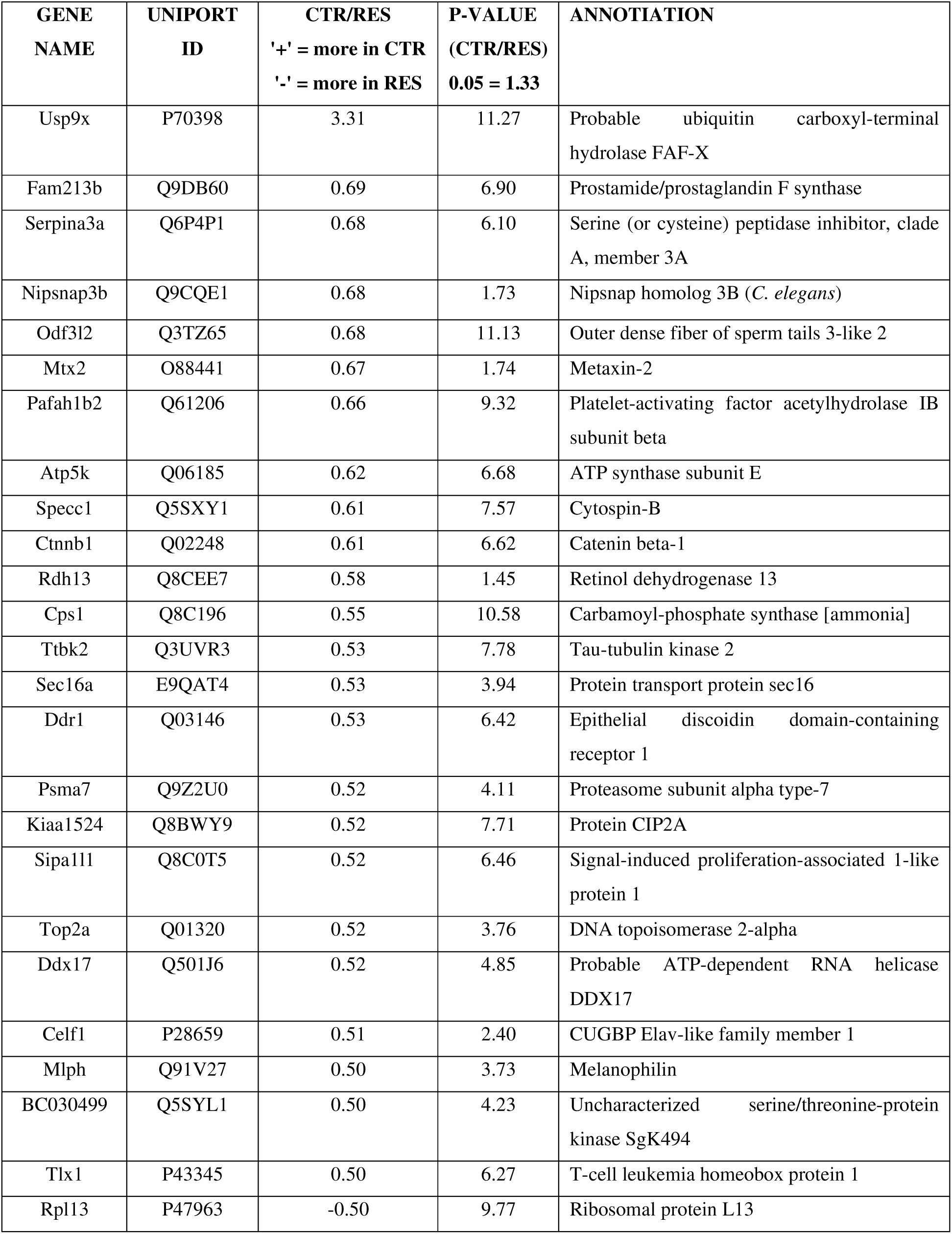
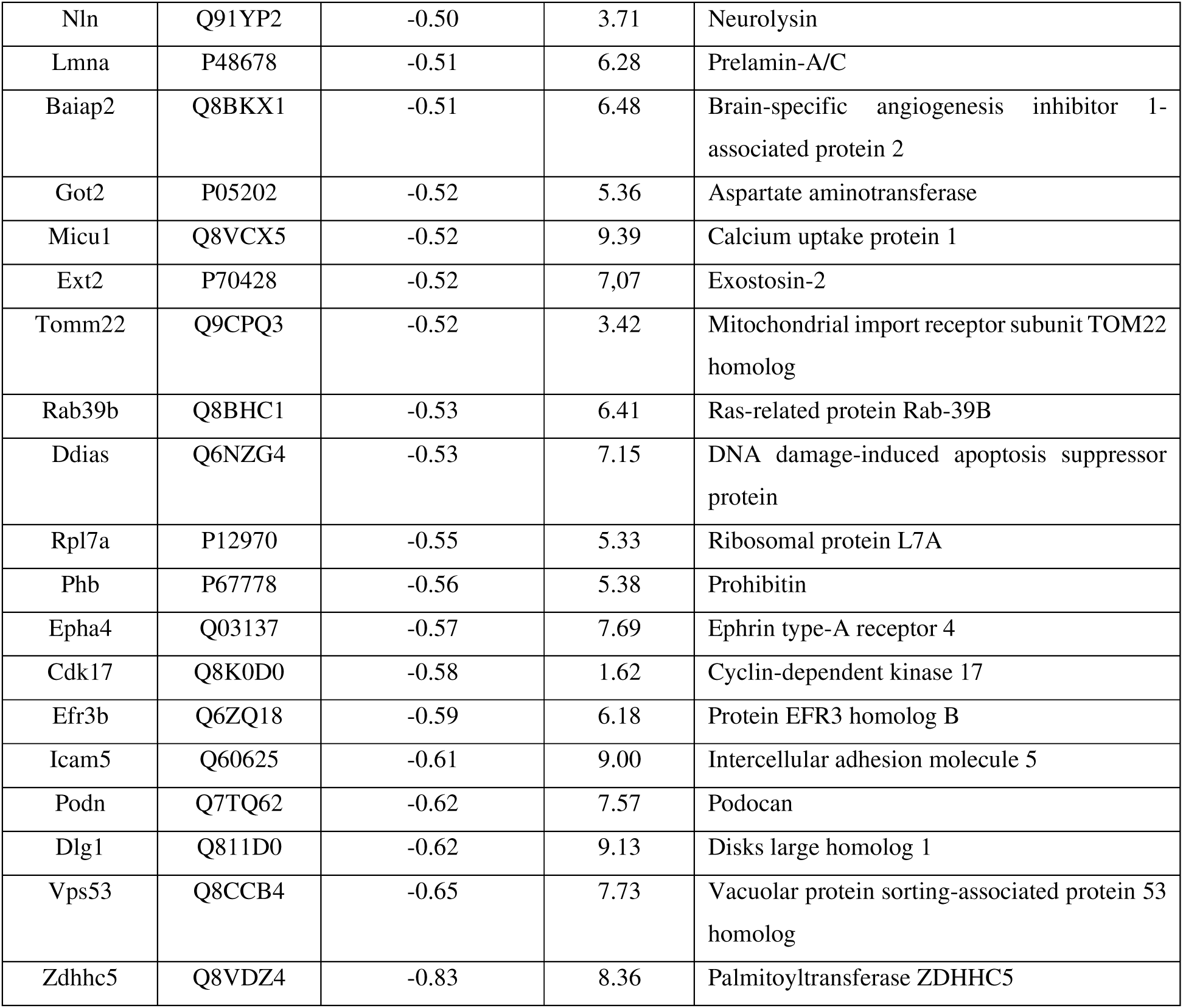

**Supplemental Table 1B.**

| GENE NAME | UNIPROT ID | CTR/ANH ('+'=more in CTR, '-'= more in ANH) | P-VALUE (CTR/ANH) 0.05 = 1.33 | ANNOTATION |
| --- | --- | --- | --- | --- |
| Usp9x | P70398 | 3.50 | 11.28 | Probable ubiquitin carboxyl-terminal hydrolase FAF-X |
| Ubr2 | Q6WKZ8 | 1.24 | 1.95 | E3 ubiquitin-protein ligase UBR2 |
| Cox4i1 | P19783 | 1.14 | 1.97 | Cytochrome c oxidase subunit 4 isoform 1 |
| Ppid | Q9CR16 | 1.08 | 1.98 | Peptidyl-prolyl cis-trans isomerase D |
| Acot10 | Q32MW3 | 1.08 | 1.98 | Acyl-coenzyme A thioesterase 10 |
| Adsl | P54822 | 0.83 | 2.30 | Adenylosuccinate lyase |
| Ppp3ca | P63328 | 0.80 | 9.94 | Serine/threonine-protein phosphatase 2B catalytic subunit alpha isoform |
| Odf3l2 | Q3TZ65 | 0.61 | 10.88 | Outer dense fiber of sperm tails 3-like 2 |
| Bcl3 | Q9Z2F6 | 0.60 | 10.96 | B-cell lymphoma 3 protein homolog |
| Maob | Q8BW75 | 0.60 | 10.90 | Amine oxidase [flavin-containing] B |
| Slc22a2<br>1 | Q9WTN6 | 0.57 | 6.97 | Solute carrier family 22 member 21 |
| Fam83d | Q9D7I8 | 0.53 | 9.62 | Protein FAM83D |
| Mtm1 | Q9Z2C5 | 0.52 | 3.35 | Myotubularin |
| Tnfaip1 | O70479 | 0.52 | 10.67 | BTB/POZ domain-containing adapter for CUL3-mediated RhoA degradation protein 2 |
| Prkg2 | Q61410 | 0.51 | 2.96 | cGMP-dependent protein kinase 2 |
| Serpina3<br>a | Q6P4P1 | 0.51 | 2.14 | Serine (or cysteine) peptidase inhibitor, clade A, member 3A |
| Maoa | Q64133 | 0.50 | 10.42 | Amine oxidase [flavin-containing] A |
| Klc3 | Q91W40 | 0.50 | 8.91 | Kinesin light chain 3 |
| Shank3 | Q4ACU6 | 0.50 | 8.75 | SH3 and multiple ankyrin repeat domains protein 3 |
| Hras | Q61411 | 0.50 | 6.95 | GTPase Hras |
| Wfs1 | P56695 | -0.50 | 4.62 | Wolframin |
| Tenm1 | Q9WTS4 | -0.51 | 3.44 | Teneurin-1 |
| Taf7l | Q9D3R9 | -0.51 | 10.40 | Transcription initiation factor TFIID subunit 7-like |
| Ext2 | P70428 | -0.52 | 12.01 | Exostosin-2 |
| Phb | P67778 | -0.52 | 11.97 | Prohibitin |
| Snx4 | Q91YJ2 | -0.52 | 3.68 | Sorting nexin-4 |
| Oxct1 | Q9D0K2 | -0.53 | 12.24 | Succinyl-CoA:3-ketoacid coenzyme A transferase 1 |
| Nln | Q91YP2 | -0.53 | 12.21 | Neurolysin |
| Epha4 | Q03137 | -0.53 | 6.97 | Ephrin type-A receptor 4 |
| Ddias | Q6NZG4 | -0.53 | 12.13 | DNA damage-induced apoptosis suppressor protein |
| Tagln3 | Q9R1Q8 | -0.54 | 12.32 | Transgelin-3 |
| Zdhhc13 | Q9CWU2 | -0.55 | 2.76 | Palmitoyltransferase ZDHHC13 |
| Efr3b | Q6ZQ18 | -0.56 | 12.38 | Protein EFR3 homolog B |
| Dlg1 | Q811D0 | -0.57 | 12.55 | Disks large homolog 1 |
| Gphn | Q8BUV3 | -0.60 | 12.64 | Gephyrin |
| Podn | Q7TQ62 | -0.61 | 12.76 | Podocan |
| Nrxn3 | Q6P9K9 | -0.63 | 1.34 | Neurexin-3 |
| Vps53 | Q8CCB4 | -0.65 | 13.01 | Vacuolar protein sorting-associated protein 53 homolog |
| Zdhhc5 | Q8VDZ4 | -0.80 | 9.94 | Palmitoyltransferase ZDHHC5 |

**Supplemental Table 1C.**

| GENE NAME | UNIPROT ID | RES/ANH<br>( '+'=more in RES,<br>'-'= more in ANH) | P-VALUE<br>(RES/ANH)<br>0.05 = 1.33 | ANNOTATION |
| --- | --- | --- | --- | --- |
| Ppp3ca | P63328 | 0.83 | 8.36 | Serine/threonine-protein phosphatase 2B catalytic subunit alpha isoform |
| Recql | Q9Z129 | -0.50 | 2.54 | ATP-dependent DNA helicase Q1 |
| Tenm1 | Q9WTS4 | -0.50 | 3.19 | Teneurin-1 |
| Fastkd1 | Q6DI86 | -0.51 | 9.54 | FAST kinase domain-containing protein 1 |
| Gcc2 | Q8CHG3 | -0.53 | 4.39 | GRIP and coiled-coil domain-containing protein 2 |
| Zdhhc13 | Q9CWU2 | -0.60 | 2.92 | Palmitoyltransferase ZDHHC13 |

**Supplemental Table 2A.**
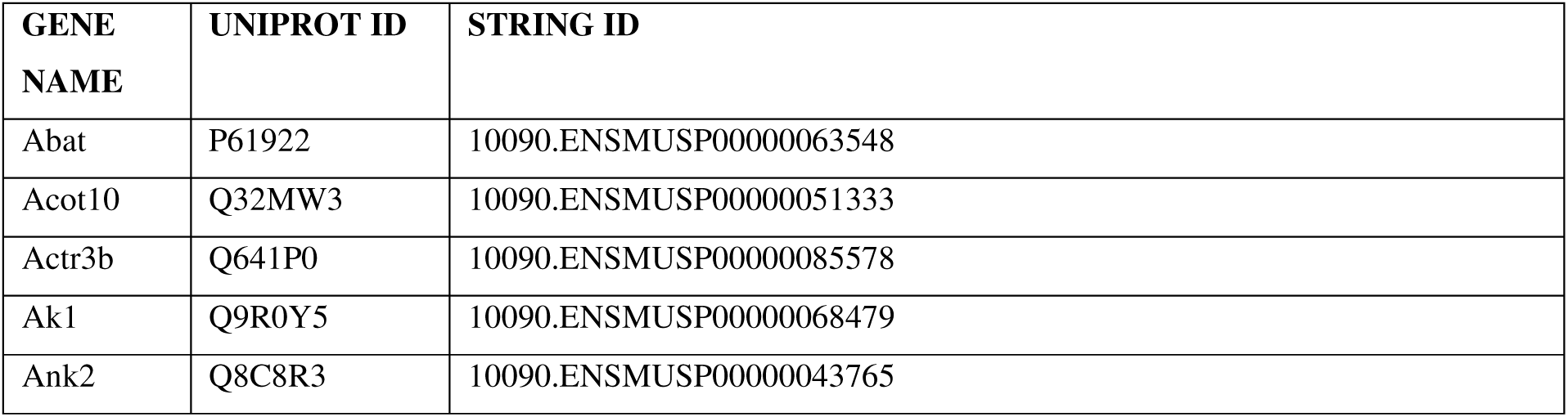
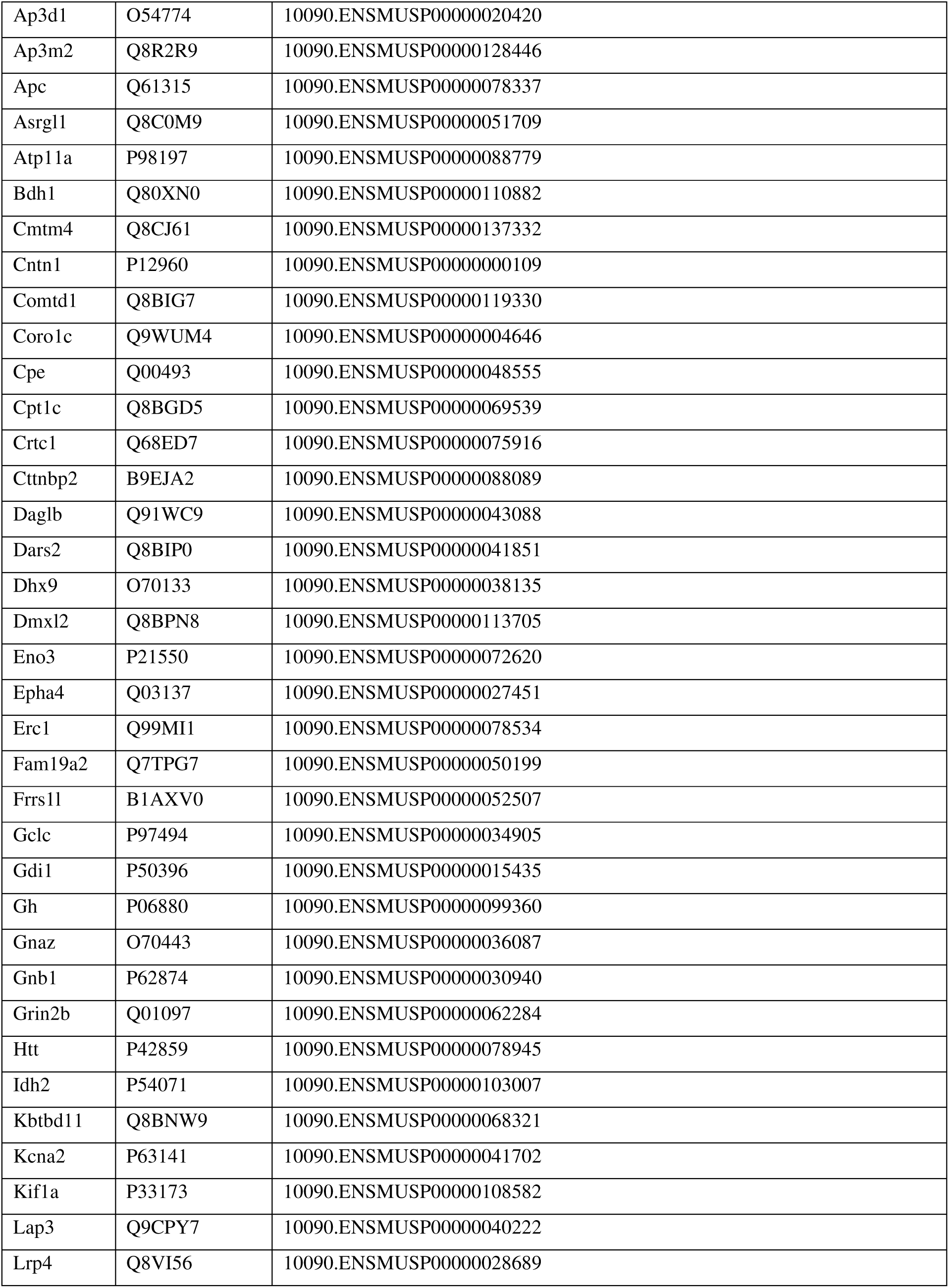
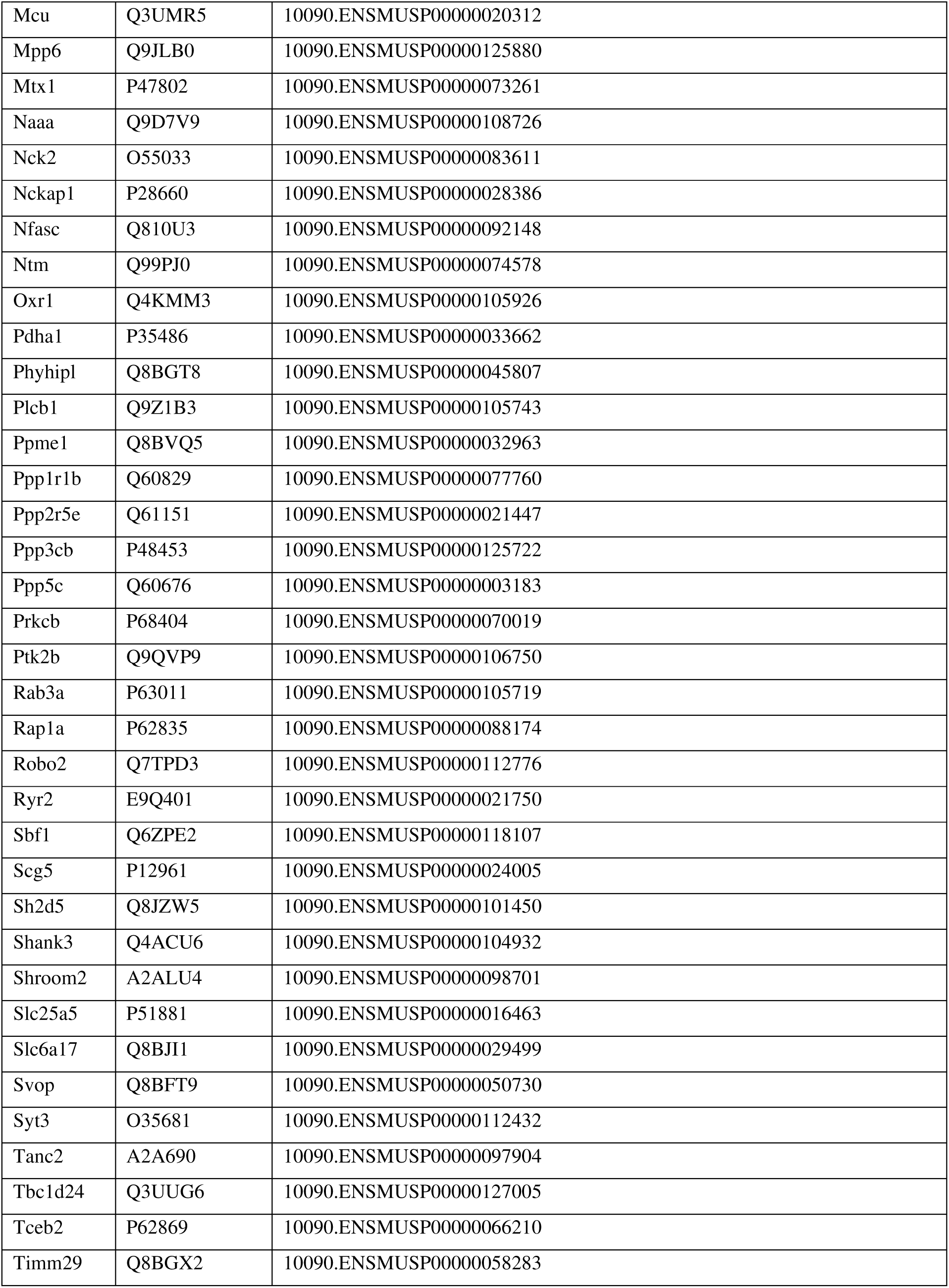
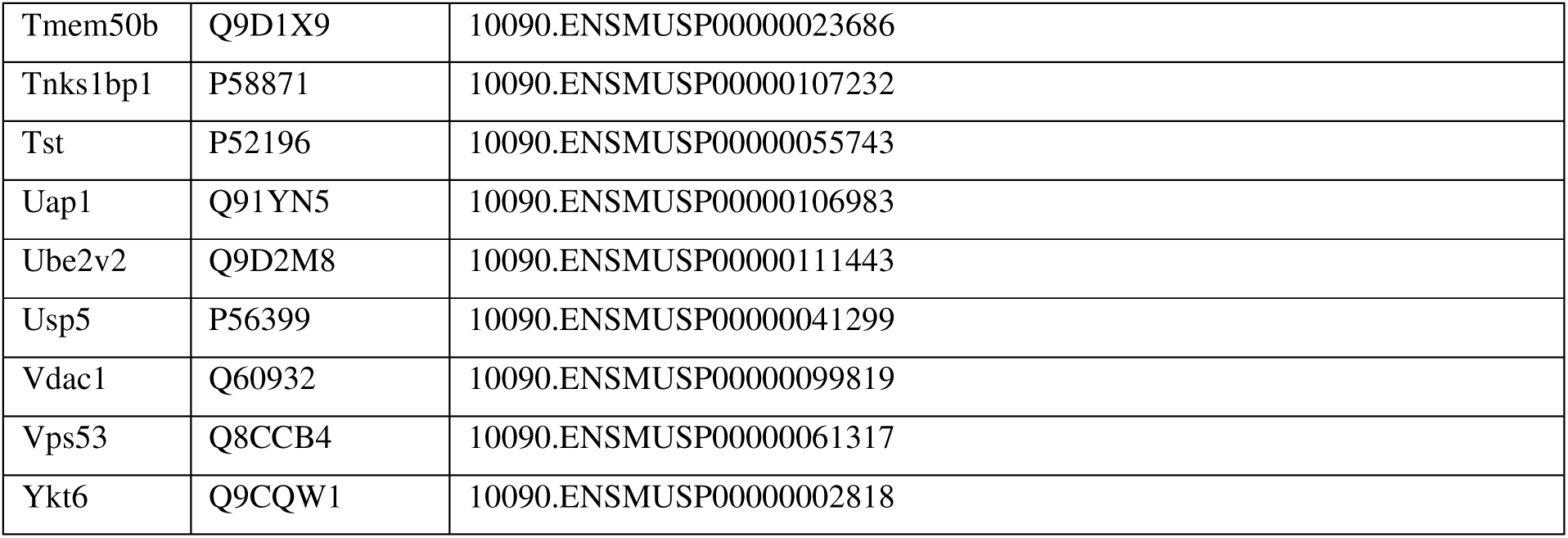

## Notes

### Competing Interest Statement

The authors have declared no competing interest.

### Summary of Updates

The revised version of the manuscript additionally includes: analysis of total palmitoylation in the CA1 region of the hippocampus (Figure 3G); analysis of spine morphology in hippocampal subregions (Figure 5); functional analysis in the CA3 region of the hippocampus.

## References

1. Suresh, S. Fatigue of Materials. (Cambridge University Press, 1998).

2. Kakani, S. L. Material Science. (New Age International (P) Ltd., Publishers, 2004).

3. Hassler, U. & Kohler, N. Resilience in the built environment. Build. Res. Inf. 42, 119–129 (2014).

4. Weichselgartner, J. & Kelman, I. Geographies of resilience: Challenges and opportunities of a descriptive concept. Prog. Hum. Geogr. 39, 249–267 (2015).

5. Rankenberg, T. et al. Age-Dependent Abiotic Stress Resilience in Plants. Trends Plant Sci. 26, 692–705 (2021).

6. Pushpalal, D. & Suzuki, A. A New Methodology for Measuring Tsunami Resilience Using Theory of Springs. Geosciences 10, 469 (2020).

7. Adger, W. N. Social and ecological resilience: are they related? Prog. Hum. Geogr. 24, 347– 364 (2000).

8. Bryan, C., O’Shea, D. & MacIntyre, T. Stressing the relevance of resilience: A systematic review of resilience across the domains of sport and work. Int. Rev. Sport Exerc. Psychol. 12, 70–111 (2019).

9. Plummer, R. & Armitage, D. A resilience-based framework for evaluating adaptive co-management: Linking ecology, economics and society in a complex world. Ecol. Econ. 61, 62–74 (2007).

10. Richter-Levin, G. & Sandi, C. Title: “Labels Matter: Is it stress or is it Trauma?” Transl. Psychiatry 11, 1–9 (2021).

11. Stern, Y. et al. Whitepaper: Defining and investigating cognitive reserve, brain reserve, and brain maintenance. Alzheimers Dement. 16, 1305–1311 (2020).

12. Richardson, G. E. The metatheory of resilience and resiliency. J. Clin. Psychol. 58, 307–321 (2002).

13. Den Hartigh, R. J. R. & Hill, Y. Conceptualizing and measuring psychological resilience: What can we learn from physics? New Ideas Psychol. 66, 100934 (2022).

14. Fletcher, D. & Sarkar, M. Psychological Resilience. Eur. Psychol. 18, 12–23 (2013).

15. Southwick, S. M., Bonanno, G. A., Masten, A. S., Panter-Brick, C. & Yehuda, R. Resilience definitions, theory, and challenges: interdisciplinary perspectives. Eur. J. Psychotraumatology 5, (2014).

16. Arenaza-Urquijo, E. M. & Vemuri, P. Resistance vs resilience to Alzheimer disease. Neurology 90, 695–703 (2018).

17. Hess, J. L. et al. A polygenic resilience score moderates the genetic risk for schizophrenia. Mol. Psychiatry 26, 800–815 (2021).

18. Davydov, D. M., Stewart, R., Ritchie, K. & Chaudieu, I. Resilience and mental health. Clin. Psychol. Rev. 30, 479–495 (2010).

19. Kane, C. et al. Clinical factors influencing resilience in patients with anorexia nervosa. Neuropsychiatr. Dis. Treat. 15, 391–395 (2019).

20. Y, M., F, W. & B, F.-A. Resilience research in schizophrenia: a review of recent developments. Curr. Opin. Psychiatry 29, (2016).

21. Wu, G. et al. Understanding resilience. Front. Behav. Neurosci. 7, 10 (2013).

22. Peña, C. J., Nestler, E. J. & Bagot, R. C. Environmental Programming of Susceptibility and Resilience to Stress in Adulthood in Male Mice. Front. Behav. Neurosci. 13, (2019).

23. Krishnan, V. et al. Molecular adaptations underlying susceptibility and resistance to social defeat in brain reward regions. Cell 131, 391–404 (2007).

24. Isingrini, E. et al. Resilience to chronic stress is mediated by noradrenergic regulation of dopamine neurons. Nat. Neurosci. 19, 560–563 (2016).

25. Cathomas, F., Murrough, J. W., Nestler, E. J., Han, M.-H. & Russo, S. J. Neurobiology of Resilience: Interface Between Mind and Body. Biol. Psychiatry 86, 410–420 (2019).

26. Highland, J. N., Zanos, P., Georgiou, P. & Gould, T. D. Group II metabotropic glutamate receptor blockade promotes stress resilience in mice. Neuropsychopharmacology 44, 1788– 1796 (2019).

27. Friedman, A. K. et al. KCNQ channel openers reverse depressive symptoms via an active resilience mechanism. Nat. Commun. 7, 11671 (2016).

28. Stainton, A. et al. Resilience as a multimodal dynamic process. Early Interv. Psychiatry 13, 725–732 (2019).

29. Krzystyniak, A. et al. Prophylactic Ketamine Treatment Promotes Resilience to Chronic Stress and Accelerates Recovery: Correlation with Changes in Synaptic Plasticity in the CA3 Subregion of the Hippocampus. Int. J. Mol. Sci. 20, (2019).

30. Brachman, R. A. et al. Ketamine as a Prophylactic Against Stress-Induced Depressive-like Behavior. Biol. Psychiatry 79, 776–786 (2016).

31. Okine, T., Shepard, R., Lemanski, E. & Coutellier, L. Sex Differences in the Sustained Effects of Ketamine on Resilience to Chronic Stress. Front. Behav. Neurosci. 14, (2020).

32. Camargo, A., Dalmagro, A. P., Wolin, I. A. V., Kaster, M. P. & Rodrigues, A. L. S. The resilient phenotype elicited by ketamine against inflammatory stressors-induced depressive-like behavior is associated with NLRP3-driven signaling pathway. J. Psychiatr. Res. 144, 118–128 (2021).

33. Becker, M., Pinhasov, A. & Ornoy, A. Animal Models of Depression: What Can They Teach Us about the Human Disease? Diagnostics 11, 123 (2021).

34. Ménard, C., Hodes, G. E. & Russo, S. J. Pathogenesis of depression: Insights from human and rodent studies. Neuroscience 321, 138–162 (2016).

35. Shadrina, M., Bondarenko, E. A. & Slominsky, P. A. Genetics Factors in Major Depression Disease. Front. Psychiatry 9, 334 (2018).

36. Malhi, G. S. & Mann, J. J. Depression. The Lancet 392, 2299–2312 (2018).

37. Schmitt, A., Malchow, B., Hasan, A. & Falkai, P. The impact of environmental factors in severe psychiatric disorders. Front. Neurosci. 8, (2014).

38. Duman, R. S., Aghajanian, G. K., Sanacora, G. & Krystal, J. H. Synaptic plasticity and depression: new insights from stress and rapid-acting antidepressants. Nat. Med. 22, 238–249 (2016).

39. Logan, R. W. et al. Chronic Stress Induces Brain Region-Specific Alterations of Molecular Rhythms that Correlate with Depression-like Behavior in Mice. Biol. Psychiatry 78, 249–258 (2015).

40. Bremner, J. D. Stress and Brain Atrophy. CNS Neurol. Disord. Drug Targets 5, 503–512 (2006).

41. Castrén, E. & Monteggia, L. M. Brain-Derived Neurotrophic Factor Signaling in Depression and Antidepressant Action. Biol. Psychiatry 90, 128–136 (2021).

42. MacQueen, G. & Frodl, T. The hippocampus in major depression: evidence for the convergence of the bench and bedside in psychiatric research? Mol. Psychiatry 16, 252–264 (2011).

43. Filatova, E. V., Shadrina, M. I. & Slominsky, P. A. Major Depression: One Brain, One Disease, One Set of Intertwined Processes. Cells 10, 1283 (2021).

44. Hasler, G. Pathophysiology of depression: do we have any solid evidence of interest to clinicians? World Psychiatry Off. J. World Psychiatr. Assoc. WPA 9, 155–161 (2010).

45. Holmes, S. E. et al. Lower synaptic density is associated with depression severity and network alterations. Nat. Commun. 10, 1529 (2019).

46. Marsden, W. N. Synaptic plasticity in depression: Molecular, cellular and functional correlates. Prog. Neuropsychopharmacol. Biol. Psychiatry 43, 168–184 (2013).

47. Price, R. B. & Duman, R. Neuroplasticity in cognitive and psychological mechanisms of depression: an integrative model. Mol. Psychiatry 25, 530–543 (2020).

48. Thompson, S. M. et al. An excitatory synapse hypothesis of depression. Trends Neurosci. 38, 279–294 (2015).

49. Autry, A. E. et al. NMDA receptor blockade at rest triggers rapid behavioural antidepressant responses. Nature 475, 91–95 (2011).

50. Duman, R. S., Li, N., Liu, R.-J., Duric, V. & Aghajanian, G. Signaling pathways underlying the rapid antidepressant actions of ketamine. Neuropharmacology 62, 35–41 (2012).

51. Treccani, G. et al. S-Ketamine Reverses Hippocampal Dendritic Spine Deficits in Flinders Sensitive Line Rats Within 1 h of Administration. Mol. Neurobiol. 56, 7368–7379 (2019).

52. Beyeler, A. Do antidepressants restore lost synapses? Science 364, 129–130 (2019).

53. Sala, C. & Segal, M. Dendritic Spines: The Locus of Structural and Functional Plasticity. Physiol. Rev. 94, 141–188 (2014).

54. Tønnesen, J. & Nägerl, U. V. Dendritic Spines as Tunable Regulators of Synaptic Signals. Front. Psychiatry 7, (2016).

55. Penzes, P., Cahill, M. E., Jones, K. A., VanLeeuwen, J.-E. & Woolfrey, K. M. Dendritic spine pathology in neuropsychiatric disorders. Nat. Neurosci. 14, 285–293 (2011).

56. Chidambaram, S. B. et al. Dendritic spines: Revisiting the physiological role. Prog. Neuropsychopharmacol. Biol. Psychiatry 92, 161–193 (2019).

57. Bączyńska, E., Pels, K. K., Basu, S., Włodarczyk, J. & Ruszczycki, B. Quantification of Dendritic Spines Remodeling under Physiological Stimuli and in Pathological Conditions. Int. J. Mol. Sci. 22, 4053 (2021).

58. Qiao, H. et al. Dendritic Spines in Depression: What We Learned from Animal Models. Neural Plast. 2016, 8056370 (2016).

59. Moda-Sava, R. N. et al. Sustained rescue of prefrontal circuit dysfunction by antidepressant-induced spine formation. Science 364, (2019).

60. Zaręba-Kozioł, M., Figiel, I., Bartkowiak-Kaczmarek, A. & Włodarczyk, J. Insights Into Protein S-Palmitoylation in Synaptic Plasticity and Neurological Disorders: Potential and Limitations of Methods for Detection and Analysis. Front. Mol. Neurosci. 11, 175 (2018).

61. Ji, B. & Skup, M. Roles of palmitoylation in structural long-term synaptic plasticity. Mol. Brain 14, 8 (2021).

62. 62. Piguel, N. H., et al. Ankyrin-G-190 palmitoylation mediates dendrite and spine morphogenesis and is altered in response to lithium. *bioRxiv* 620708 (2019) doi:10.1101/620708.

63. Mumby, S. M. Reversible palmitoylation of signaling proteins. Curr. Opin. Cell Biol. 9, 148– 154 (1997).

64. Milligan, G., Parenti, M. & Magee, A. I. The dynamic role of palmitoylation in signal transduction. Trends Biochem. Sci. 20, 181–187 (1995).

65. Fukata, Y. & Fukata, M. Protein palmitoylation in neuronal development and synaptic plasticity. Nat. Rev. Neurosci. 11, 161–175 (2010).

66. Kang, R. et al. Neural Palmitoyl-Proteomics Reveals Dynamic Synaptic Palmitoylation. Nature 456, 904–909 (2008).

67. Jeyifous, O., et al. Palmitoylation regulates glutamate receptor distributions in postsynaptic densities through control of PSD95 conformation and orientation. Proc. Natl. Acad. Sci. 113, E8482–E8491 (2016).

68. Bijata, M. et al. Activation of the 5-HT7 receptor and MMP-9 signaling module in the hippocampal CA1 region is necessary for the development of depressive-like behavior. Cell Rep. 38, 110532 (2022).

69. Bijata, M., Bączyńska, E. & Wlodarczyk, J. A chronic unpredictable stress protocol to model anhedonic and resilient behaviors in C57BL/6J mice. STAR Protoc. 3, 101659 (2022).

70. Retana-Márquez, S. et al. Body weight gain and diurnal differences of corticosterone changes in response to acute and chronic stress in rats. Psychoneuroendocrinology 28, 207–227 (2003).

71. Tamashiro, K. L., Sakai, R. R., Shively, C. A., Karatsoreos, I. N. & Reagan, L. P. Chronic stress, metabolism, and metabolic syndrome. Stress 14, 468–474 (2011).

72. Zareba-Koziol, M. et al. Stress-induced Changes in the S-palmitoylation and S-nitrosylation of Synaptic Proteins. Mol. Cell. Proteomics MCP 18, 1916–1938 (2019).

73. Zaręba-Kozioł, M., Szwajda, A., Dadlez, M., Wysłouch-Cieszyńska, A. & Lalowski, M. Global analysis of S-nitrosylation sites in the wild type (APP) transgenic mouse brain-clues for synaptic pathology. Mol. Cell. Proteomics MCP 13, 2288–2305 (2014).

74. Gorinski, N. et al. Attenuated palmitoylation of serotonin receptor 5-HT1A affects receptor function and contributes to depression-like behaviors. Nat. Commun. 10, 1–14 (2019).

75. Brzdąk, P., Włodarczyk, J., Mozrzymas, J. W. & Wójtowicz, T. Matrix Metalloprotease 3 Activity Supports Hippocampal EPSP-to-Spike Plasticity Following Patterned Neuronal Activity via the Regulation of NMDAR Function and Calcium Flux. Mol. Neurobiol. 54, 804–816 (2017).

76. Wójtowicz, T. & Mozrzymas, J. W. Matrix metalloprotease activity shapes the magnitude of EPSPs and spike plasticity within the hippocampal CA3 network. Hippocampus 26, 414 (2016).

77. Kim, E., Owen, B., Holmes, W. R. & Grover, L. M. Decreased afferent excitability contributes to synaptic depression during high-frequency stimulation in hippocampal area CA1. J. Neurophysiol. 108, 1965–1976 (2012).

78. Brzdak, P. et al. Synaptic Potentiation at Basal and Apical Dendrites of Hippocampal Pyramidal Neurons Involves Activation of a Distinct Set of Extracellular and Intracellular Molecular Cues. Cereb. Cortex N. Y. N 1991 29, 283–304 (2019).

79. Richter-Levin, G. & Sandi, C. Title: “Labels Matter: Is it stress or is it Trauma?” Transl. Psychiatry 11, 1–9 (2021).

80. Pushpalal, D. & Suzuki, A. A New Methodology for Measuring Tsunami Resilience Using Theory of Springs. Geosciences 10, 469 (2020).

81. Tang, M. et al. Hippocampal proteomic changes of susceptibility and resilience to depression or anxiety in a rat model of chronic mild stress. Transl. Psychiatry 9, 1–12 (2019).

82. D, R., et al. Hippocampal extracellular matrix alterations contribute to cognitive impairment associated with a chronic depressive-like state in rats. Sci. Transl. Med. 9, (2017).

83. Han, X. et al. iTRAQ-based quantitative analysis of hippocampal postsynaptic density-associated proteins in a rat chronic mild stress model of depression. Neuroscience 298, 220– 292 (2015).

84. McEwen, B. S. et al. Mechanisms of stress in the brain. Nat. Neurosci. 18, 1353–1363 (2015).

85. Perić, I., Costina, V., Stanisavljević, A., Findeisen, P. & Filipović, D. Proteomic characterization of hippocampus of chronically socially isolated rats treated with fluoxetine: Depression-like behaviour and fluoxetine mechanism of action. Neuropharmacology 135, 268–283 (2018).

86. Gottschalk, M. G. et al. Fluoxetine, not donepezil, reverses anhedonia, cognitive dysfunctions and hippocampal proteome changes during repeated social defeat exposure. Eur. Neuropsychopharmacol. J. Eur. Coll. Neuropsychopharmacol. 28, 195–210 (2018).

87. McEwen, B. S., Nasca, C. & Gray, J. D. Stress Effects on Neuronal Structure: Hippocampus, Amygdala, and Prefrontal Cortex. Neuropsychopharmacology 41, 3–23 (2016).

88. Popoli, M., Yan, Z., McEwen, B. S. & Sanacora, G. The stressed synapse: the impact of stress and glucocorticoids on glutamate transmission. Nat. Rev. Neurosci. 13, 22–37 (2012).

89. Yang, Y., Ju, W., Zhang, H. & Sun, L. Effect of Ketamine on LTP and NMDAR EPSC in Hippocampus of the Chronic Social Defeat Stress Mice Model of Depression. Front. Behav. Neurosci. 12, (2018).

90. Strekalova, T., Spanagel, R., Bartsch, D., Henn, F. A. & Gass, P. Stress-Induced Anhedonia in Mice is Associated with Deficits in Forced Swimming and Exploration. Neuropsychopharmacology 29, 2007–2017 (2004).

91. Strekalova, T. & Steinbusch, H. W. M. Measuring behavior in mice with chronic stress depression paradigm. Prog. Neuropsychopharmacol. Biol. Psychiatry 34, 348–361 (2010).

92. Parsons, S., Kruijt, A.-W. & Fox, E. A Cognitive Model of Psychological Resilience. J. Exp. Psychopathol. 7, 296–310 (2016).

93. Miyagi, T. et al. Psychological resilience is correlated with dynamic changes in functional connectivity within the default mode network during a cognitive task. Sci. Rep. 10, 17760 (2020).

94. Pfarr, J.-K. et al. Brain structural connectivity, anhedonia, and phenotypes of major depressive disorder: A structural equation model approach. Hum. Brain Mapp. 42, 5063 (2021).

95. Li, N. et al. mTOR-dependent synapse formation underlies the rapid antidepressant effects of NMDA antagonists. Science 329, 959–964 (2010).

96. Ng, L. H. L. et al. Ketamine and selective activation of parvalbumin interneurons inhibit stress-induced dendritic spine elimination. Transl. Psychiatry 8, 1–15 (2018).

97. Shao, L.-X. et al. Psilocybin induces rapid and persistent growth of dendritic spines in frontal cortex in vivo. Neuron 109, 2535–2544.e4 (2021).

98. Kim, E. J., Pellman, B. & Kim, J. J. Stress effects on the hippocampus: a critical review. Learn. Mem. 22, 411–416 (2015).

99. Goldwater, D. S. et al. Structural and functional alterations to rat medial prefrontal cortex following chronic restraint stress and recovery. Neuroscience 164, 798–808 (2009).

100. Radley, J. J. et al. Reversibility of apical dendritic retraction in the rat medial prefrontal cortex following repeated stress. Exp. Neurol. 196, 199–203 (2005).

101. Yang, C. et al. Mechanistic Target of Rapamycin–Independent Antidepressant Effects of (R)-Ketamine in a Social Defeat Stress Model. Biol. Psychiatry 83, 18–28 (2018).

102. Suo, L. et al. Predictable Chronic Mild Stress in Adolescence Increases Resilience in Adulthood. Neuropsychopharmacology 38, 1387–1400 (2013).

103. Xu, D. et al. Hippocampal mTOR signaling is required for the antidepressant effects of paroxetine. Neuropharmacology 128, 181–195 (2018).

104. Sarbassov, D. D. et al. Rictor, a novel binding partner of mTOR, defines a rapamycin-insensitive and raptor-independent pathway that regulates the cytoskeleton. Curr. Biol. CB 14, 1296–1302 (2004).

105. McCabe, M. P. et al. Genetic inactivation of mTORC1 or mTORC2 in neurons reveals distinct functions in glutamatergic synaptic transmission. eLife 9, e51440 (2020).

106. Angliker, N. & Rüegg, M. A. In vivo evidence for mTORC2-mediated actin cytoskeleton rearrangement in neurons. BioArchitecture 3, 113–118 (2013).

107. Zhu, P. J., Chen, C.-J., Mays, J., Stoica, L. & Costa-Mattioli, M. mTORC2, but not mTORC1, is required for hippocampal mGluR-LTD and associated behaviors. Nat. Neurosci. 21, 799–802 (2018).

108. Kim, S.-G. et al. Tanc2-mediated mTOR inhibition balances mTORC1/2 signaling in the developing mouse brain and human neurons. Nat. Commun. 12, 2695 (2021).

109. Li, X. & Jope, R. S. Is glycogen synthase kinase-3 a central modulator in mood regulation? Neuropsychopharmacol. Off. Publ. Am. Coll. Neuropsychopharmacol. 35, 2143– 2154 (2010).

110. Kondratiuk, I. et al. GSK-3β and MMP-9 Cooperate in the Control of Dendritic Spine Morphology. Mol. Neurobiol. 54, 200–211 (2017).

111. Duda, P., Hajka, D., Wójcicka, O., Rakus, D. & Gizak, A. GSK3β: A Master Player in Depressive Disorder Pathogenesis and Treatment Responsiveness. Cells 9, 727 (2020).

112. Beurel, E., Grieco, S. F. & Jope, R. S. Glycogen synthase kinase-3 (GSK3): Regulation, actions, and diseases. Pharmacol. Ther. 148, 114–131 (2015).

113. Koo, J., Wu, X., Mao, Z., Khuri, F. R. & Sun, S.-Y. Rictor Undergoes Glycogen Synthase Kinase 3 (GSK3)-dependent, FBXW7-mediated Ubiquitination and Proteasomal Degradation. J. Biol. Chem. 290, 14120–14129 (2015).

114. Evangelisti, C., Chiarini, F., Paganelli, F., Marmiroli, S. & Martelli, A. M. Crosstalks of GSK3 signaling with the mTOR network and effects on targeted therapy of cancer. Biochim. Biophys. Acta BBA - Mol. Cell Res. 1867, 118635 (2020).

115. Vidal, R. et al. Targeting β-Catenin in GLAST-Expressing Cells: Impact on Anxiety and Depression-Related Behavior and Hippocampal Proliferation. Mol. Neurobiol. 56, 553–566 (2019).

116. Hayashi, T., Rumbaugh, G. & Huganir, R. L. Differential regulation of AMPA receptor subunit trafficking by palmitoylation of two distinct sites. Neuron 47, 709–723 (2005).

117. Lin, D. et al. Regulation of AMPA receptor extrasynaptic insertion by 4.1N, phosphorylation and palmitoylation. Nat. Neurosci. 12, 879–887 (2009).

118. Wei, J., Liu, W. & Yan, Z. Regulation of AMPA Receptor Trafficking and Function by Glycogen Synthase Kinase 3 *. J. Biol. Chem. 285, 26369–26376 (2010).

119. Hausser, A. & Schlett, K. Coordination of AMPA receptor trafficking by Rab GTPases. Small GTPases 10, 419–432 (2019).

120. Purkey, A. M. & Dell’Acqua, M. L. Phosphorylation-Dependent Regulation of Ca2+- Permeable AMPA Receptors During Hippocampal Synaptic Plasticity. Front. Synaptic Neurosci. 12, (2020).

121. Lee, H.-K., Takamiya, K., He, K., Song, L. & Huganir, R. L. Specific Roles of AMPA Receptor Subunit GluR1 (GluA1) Phosphorylation Sites in Regulating Synaptic Plasticity in the CA1 Region of Hippocampus. J. Neurophysiol. 103, 479–489 (2010).

122. Lu, W. et al. Phosphorylation of Tyrosine 1070 at the GluN2B Subunit Is Regulated by Synaptic Activity and Critical for Surface Expression of N-Methyl-d-aspartate (NMDA) Receptors *. J. Biol. Chem. 290, 22945–22954 (2015).

123. Ruxton, G. D. & Neuhäuser, M. Improving the reporting of P-values generated by randomization methods. Methods Ecol. Evol. 4, 1033–1036 (2013).

124. Davison, A. C. & Hinkley, D. V. Bootstrap Methods and their Application. (Cambridge University Press, 1997). doi:10.1017/CBO9780511802843.

125. Magnowska, M. et al. Transient ECM protease activity promotes synaptic plasticity. Sci. Rep. 6, 27757 (2016).

126. Degasperi, A. et al. Evaluating Strategies to Normalise Biological Replicates of Western Blot Data. PLOS ONE 9, e87293 (2014).

